# Environment-conditioned design of *α*-helical peptides

**DOI:** 10.64898/2026.05.07.723485

**Authors:** Daniel Conde-Torres, Rebeca García-Fandiño, Ángel Piñeiro

## Abstract

Designing peptide sequences that remain stable and selective across heterogeneous environments remains a central challenge in biomolecular modeling. Here we introduce an interpretable, physics-based Hamiltonian for environment-conditioned design of *α*-helical peptide sequences. The model integrates helix propensities, pairwise interactions, electrostatics, anisotropic solvent exposure, and interfacial geometry into a unified energy function. To enable comparison across sequence lengths and environments, all contributions are rescaled and expressed as Z-scores relative to random sequence ensembles, yielding a normalized design landscape with balanced physical terms. This formulation defines a structured optimization problem that can be explored using exact, heuristic, and hybrid quantum– classical approaches without modification of the underlying model. The Hamiltonian recovers polar and apolar limits, discriminates experimentally characterized water-soluble and transmembrane *α*-helical peptide sequences, and captures the preferential stabilization of membrane-active sequences at anionic interfaces over non-functional controls. It further enables multi-objective and selective design, generating candidate sequences with tunable environmental specificity.

---

Designing biomolecular sequences that remain stable and functionally compatible across different physical environments is a fundamental challenge at the interface of physics, chemistry, and computation. In many relevant systems, sequence stability is not solely determined by intrinsic folding preferences, but emerges from a context-dependent balance of interactions with the surrounding medium, including solvent polarity, electrostatics, and interfacial geometry [1, 2]. From a computational perspective, this defines a high-dimensional optimization problem in which sequences correspond to discrete configurations evaluated under environment-dependent energetic constraints.

A particularly illustrative case is provided by membrane-interacting *α*-helical peptides, in particular antimicrobial peptides (AMPs), whose function depends on their ability to selectively stabilize amphipathic conformations at heterogeneous interfaces. In many therapeutically relevant settings, such as antimicrobial activity, these peptides do not act by binding to a specific protein target but by recognizing and disrupting particular membrane environments [3, 4]. Their activity is therefore governed by preferential compatibility with anionic bacterial or diseased-cell membranes relative to the more neutral membranes of healthy mammalian cells [4, 5]. This context-dependent selectivity makes membrane-active *α*-helical peptides a useful model system for studying environment-conditioned sequence design [6, 7, 8].

Modern structure prediction and design methods have been transformed by deep-learning models such as AlphaFold 3 [9] and related architectures [10]. These approaches are powerful, but they typically rely on learned statistical correlations, homology information or large training corpora, and their scores do not always expose how solvent polarity, charge and mem-brane geometry compete in a new design setting [11, 12, 13]. Physics-based approaches make these interactions explicit, but exhaustive exploration of sequence–conformation space becomes rapidly intractable [14, 15, 16].

Here a physics-based Hamiltonian framework is introduced for sequence design on membrane-interfacing *α*-helical scaffolds. The design problem is mapped onto an energy function related to generalized Potts and spin-glass models [17, 18, 19], combining helix propensities, local sequence context, residue contacts, solvent polarity, membrane charge coupling and electrostatic screening [20, 21]. Sequence generation is therefore driven by an explicit energy landscape: low-energy sequences are favored because they are compatible with a prescribed helical state in a specified environment, not because they resemble sequences learned from a training set.

The same normalized Hamiltonian can be inspected term by term, compared across environments through random-reference Z-scores, sampled as an energy landscape and optimized under alternative objectives. It can also be explored with exact enumeration for small systems, classical heuristics for larger systems and hybrid quantum-classical workflows such as QAOA without changing the physical objective. The model is validated through reference-sequence recognition against experimentally characterized *α*-helical peptide systems, explicit negative controls and landscape diagnostics. A web implementation is available at https://simbios.usc.es/HelixDesignStudio/.

## 1 Results

A peptide sequence is evaluated on a fixed helical scaffold with an environment-dependent exposure pattern; the resulting energy terms are standardized against random sequence backgrounds to obtain comparable Z-scores; and the same scoring model is then used for reference recognition, prospective design and landscape diagnostics (Figure 1; see Methods). Random-reference Z-scores place different lengths, alphabets and environments on a common scale. Reference-sequence benchmarks test measured systems outside the optimization set, whereas negative controls and landscape diagnostics test whether favorable scores reflect membrane-active organization instead of generic helicity.

**Figure 1:**
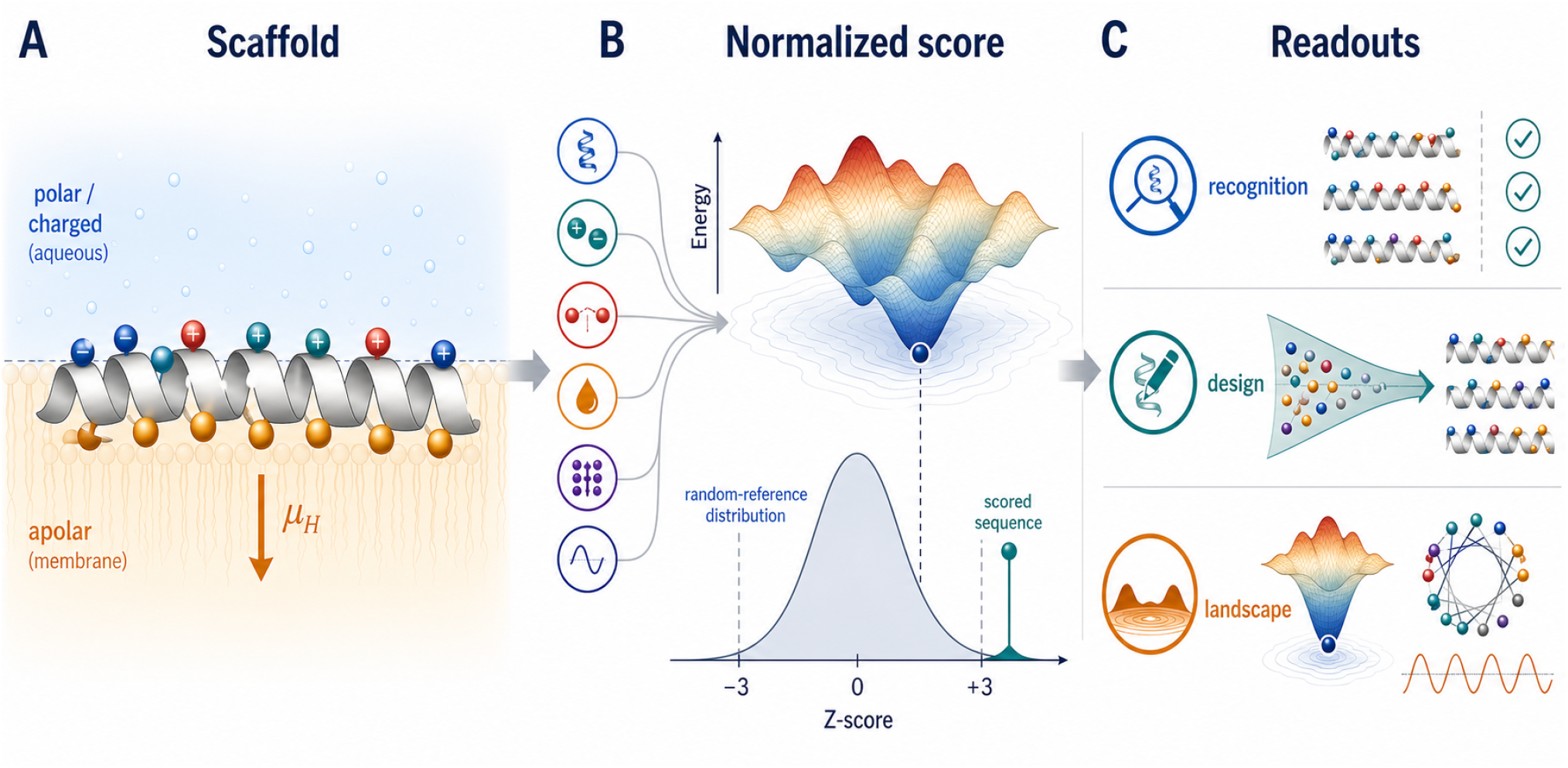
Environment-conditioned Hamiltonian used for helical peptide scoring and design. **(A)** A membrane-parallel amphipathic helix is evaluated at an interface, with hydrophobic residues oriented toward the apolar phase and polar or charged residues toward the aqueous phase. **(B)** Physical energy terms are combined into a normalized Hamiltonian score and compared against random-reference backgrounds to obtain environment-specific Z-scores. **(C)** The same scoring model is used for reference recognition, prospective design and sequence-landscape diagnostics.

### 1.1 Recognition of Water-Soluble and Transmembrane Model Peptides

External recognition was first assessed by evaluating known aqueous alanine-rich Glu/Lys peptides and hydrophobic WALP/GWALP transmembrane model peptides in both homogeneous polar and homogeneous apolar Hamiltonians (Table 1). These reference systems have experimentally established helical behavior and were not generated by the optimizer.

**Table 1:**
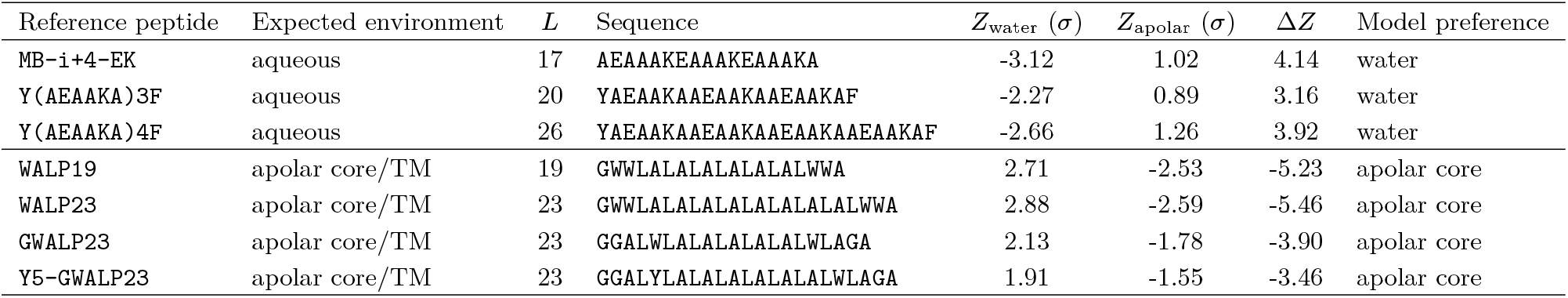
Recognition of experimentally characterized water-soluble and transmembrane model peptides. Aqueous references are alanine-rich Glu/Lys peptides from short water-soluble helix studies; apolar-core references are WALP/GWALP transmembrane model peptides. *Z*_water_ and *Z*_apolar_ are homogeneous-medium Z-scores in units of *σ* relative to 10,000 random sequences of the same length and alphabet. Terminal acetylation, amidation, or ethanolamide groups present in some experimental references are not represented in the sequence-only Hamiltonian. Δ*Z* = *Z*_apolar_ *− Z*_water_; positive values indicate water recognition, whereas negative values indicate apolar-core recognition.

The sign separation is clear. The aqueous reference peptides have favorable water Z-scores and unfavorable apolar Z-scores, with Δ*Z* = *Z*_apolar_ − *Z*_water_ between 3.16 and 4.14. Conversely, WALP and GWALP reference peptides have unfavorable water Z-scores and favorable apolar Z-scores, with Δ*Z* between −3.46 and −5.46. Thus, using only the sign of Δ*Z* as a classifier, all reference sequences are assigned to the expected environmental class. These results support the physical interpretation of the homogeneous polar/apolar scoring terms; the apolar references correspond to transmembrane apolar-core model peptides, used here as model systems for low-polarity helical compatibility.

### 1.2 Recognition of Canonical Membrane-Active Peptides

The interfacial Hamiltonian was benchmarked against membrane-active reference peptides whose amphipathic *α*-helical behavior at negatively charged interfaces is experimentally established. Seven membrane-active peptides (`LL-37`, `Melittin`, `Magainin 2`, `Cecropin A`, `PGLa`, `Piscidin 1` and `BP100`) were evaluated together with two non-AMP helix-forming controls (`(EAAAK)3` and `Poly-Ala 15`). Each sequence was aligned by its transverse hydrophobic moment and scanned over anionic, neutral and cationic interfaces with sampled penetrations from 5% to 95%. Z-scores were computed relative to 3,000 random sequences of the same length over the full 20-residue alphabet.

All seven AMPs are recognized as more favorable in an anionic interface than in homogeneous water or homogeneous apolar medium, with best anionic Z-scores from −1.71 to −5.56 (Figure 2). In every AMP case, the anionic optimum is also more favorable than the neutral or cationic interfacial optimum. The two non-AMP controls show the opposite behavior: `(EAAAK)3` and `Poly-Ala 15` remain more favorable in water than in the anionic interface. The benchmark therefore separates established anionic-membrane-active peptides from generic helix-forming controls. The complete numerical table is provided in the Supplementary Information.

**Figure 2:**
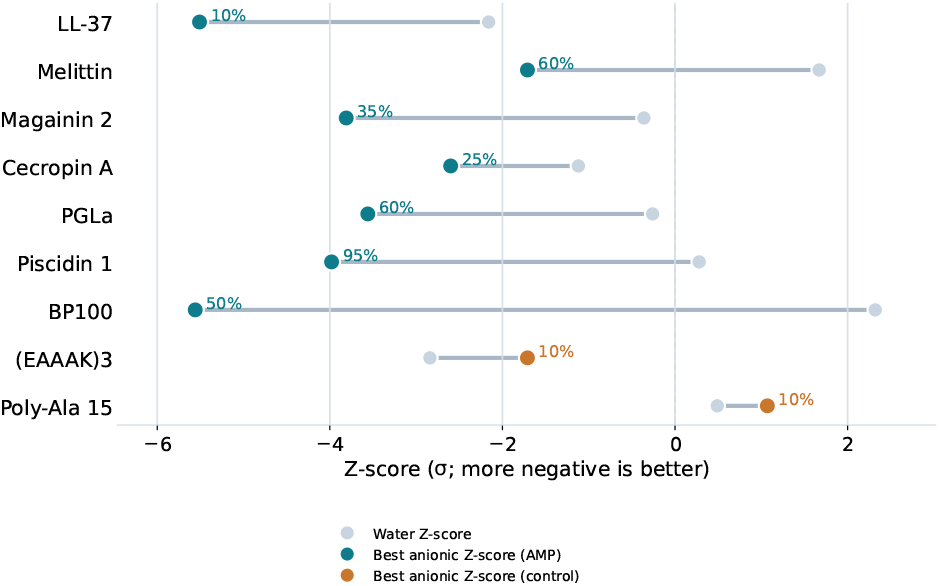
Recognition of membrane-active benchmark peptides by the anionic interfacial Hamiltonian. For each peptide, the gray marker indicates the Z-score in homogeneous water and the colored marker shows the best Z-score found in the anionic interfacial scan; both are reported in units of *σ*. Blue-green markers denote established membrane-active AMPs and brown markers denote non-AMP helix-forming controls. The numbers next to the interfacial markers indicate the optimal sampled anionic penetration. The AMP set consistently shifts toward more favorable Z-scores under anionic interfacial conditions, whereas the aqueous helix-forming controls do not.

### 1.3 Environment-Conditioned Sequence Design

The same Hamiltonian can also be used prospectively by changing the objective rather than the scoring terms. As a first test, sequences were optimized against both polar and apolar homogeneous Hamiltonians. Using the joint objective in Eq. 26, sequence-space annealing at *L* = 20 shows a transition from polar-biased optima to balanced cross-environment candidates as the gap penalty is increased. Without a gap penalty, the best sequence is strongly polar-biased, EKEKFEKEKEKEKFEKEKEK, with *Z*_polar_ = −9.26 and *Z*_apolar_ = −1.29. Introducing the balance term yields mixed aromatic/charged sequences enriched in Glu, Lys, Phe and Trp. At λ_gap_ = 0.5 and 1, both environments remain favorable, with Z-scores between approximately −3.4 and −3.8; at λ_gap_ = 4, the normalized polar–apolar energy difference is reduced to 0.02. The full sequence table is provided in the Supplementary Information.

Interfacial selectivity was then examined in a one-vs-rest design mode. For the membrane-targeting problem, the target was the anionic interface, and the off-target set comprised the neutral interface, the cationic interface, homogeneous water and homogeneous apolar medium. The interpolating objective in Eq. 27 was evaluated for λ = 0, 0.25, 0.5 and 0.75 at fixed penetrations of 10%, 25%, 50%, 75% and 90%.

Figure 3 summarizes the recovered candidates. The λ > 0 branch is stable: λ = 0.25, 0.5 and 0.75 recover the same top-ranked candidate at every fixed penetration depth. Shallow insertions remain favorable in the anionic interface but are also too favorable in water, whereas deep insertions become too compatible with homogeneous apolar medium; the 50% case gives the best compromise in this branch. The hard-selective limit λ = 0 yields a distinct family with cleaner one-vs-rest separation, especially at shallow-to-intermediate penetration. These sequences are model predictions for follow-up testing. Homogeneous boundary-case optima and representative interfacial wheel plots are provided in the Supplementary Information as sanity checks on the force balance.

**Figure 3:**
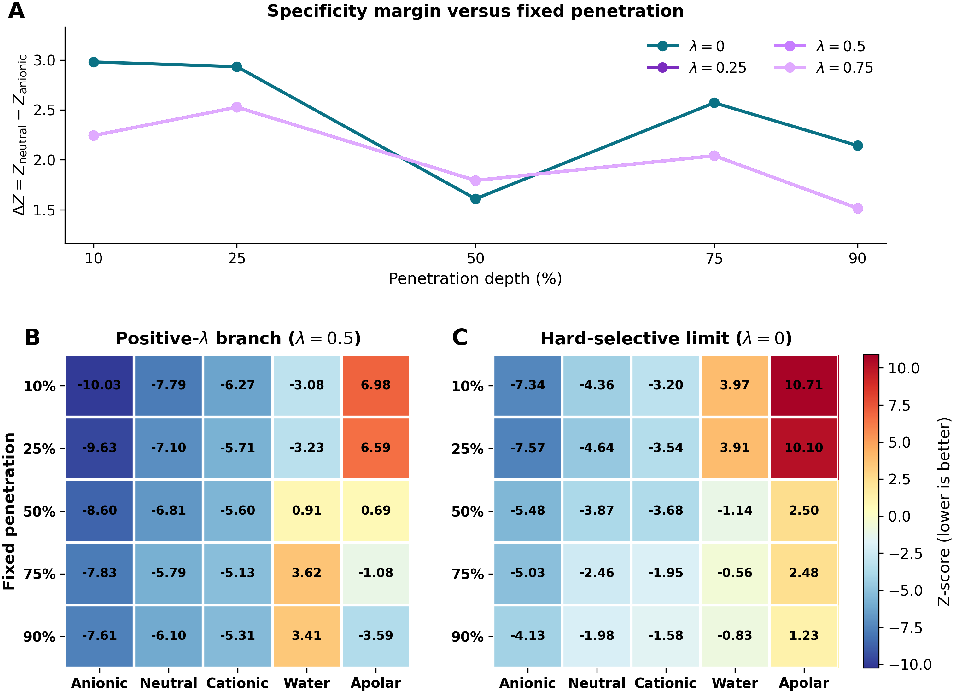
Lambda-dependent one-vs-rest specificity design for anionic membranes. Each point or heatmap row corresponds to a single top-ranked *L* = 20 sequence obtained for the indicated objective and fixed penetration depth; the corresponding full candidate libraries are provided in the Supplementary Information. The selected sequence is then rescored in the anionic, neutral, cationic, homogeneous-water, and homogeneous-apolar environments, with all Z-scores reported relative to random sequences of the same length and alphabet. **(A)** Specificity margin Δ*Z* = *Z*_neutral_ − *Z*_anionic_ of the top-ranked design at each imposed penetration depth for *λ* = 0, 0.25, 0.5, and 0.75. The curves for *λ* = 0.25, 0.5, and 0.75 overlap exactly at the level of the top-ranked design. **(B)** Z-score heatmap for the representative positive-*λ* branch (*λ* = 0.5), showing the best candidate recovered at each penetration; the same top-ranked sequence is obtained for *λ* = 0.25 and 0.75. **(C)** Z-score heatmap for the hard-selective limit *λ* = 0, which yields a distinct sequence family with stronger shallow-penetration separation from homogeneous water and apolar medium. In all cases the most competitive off-target remains the neutral interface.

### 1.4 Sequence-Landscape Structure: Funnels, Periodicity, and Neutral Networks

The interfacial sequence landscape was analyzed to test whether low-energy solutions have collective structure. For *N* = 50, 000 random *L* = 12 sequences over the reduced 16-residue alphabet, total energy was compared with amphipathicity and net charge. This analysis follows the logic of energy-landscape theory, where productive biomolecular landscapes are organized by coherent low-energy basins instead of isolated unrelated minima [19, 22].

Low-energy sequences become enriched in larger transverse hydrophobic moments even though this descriptor is used only for analysis (Figure 4A). Net charge remains constrained by the electrostatic terms (Figure 4B). After steepest-descent optimization of 1,000 random decoys, the optimized ensemble forms a separated low-energy basin with a design Z-score of −10.37 σ relative to 10,000 random sequences (Figure 4C). Its average helical wheel shows a consistent amphipathic face (Figure 4D), and the hydrophobicity autocorrelation has peaks at *k* = 3 and *k* = 4 (Figure 4E), matching the periodicity of an *α*-helix. Additional distance-to-optimum and variance diagnostics are provided in the Supplementary Information.

**Figure 4:**
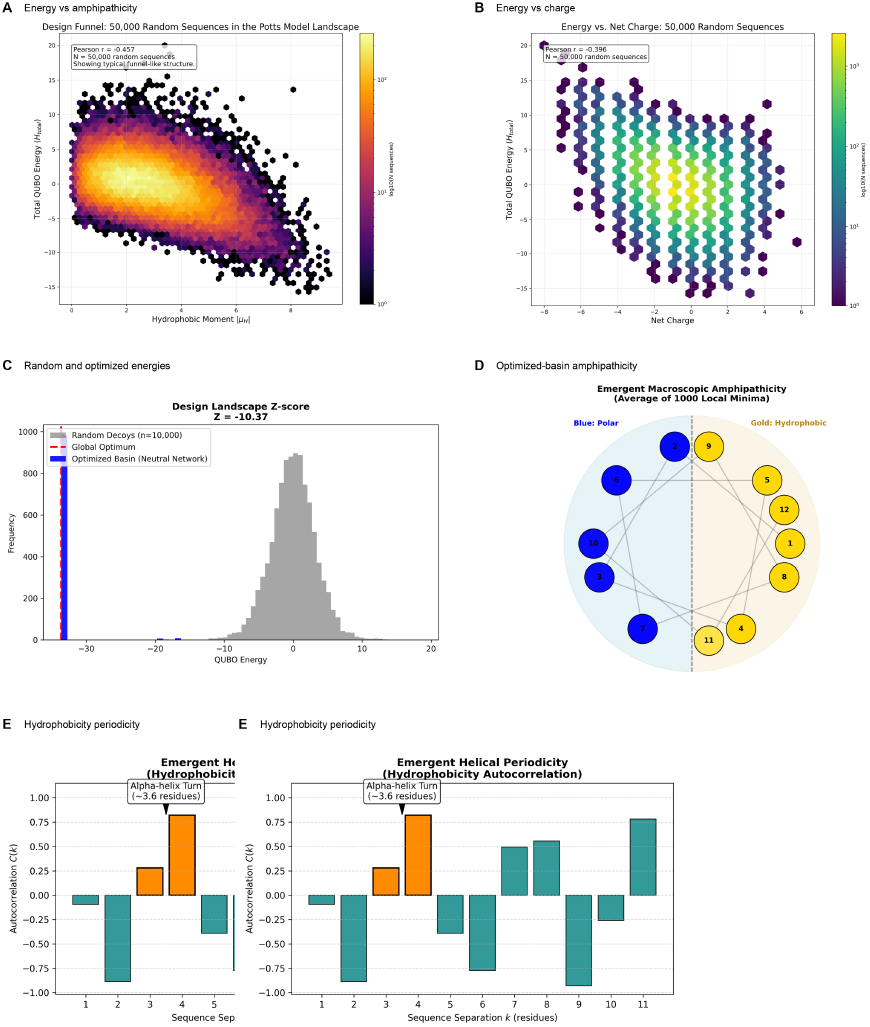
Collective structure of the generated interfacial sequence landscape. **(A)** Total Hamiltonian energy versus transverse hydrophobic moment for 50,000 random *L* = 12 sequences over the reduced 16-residue working alphabet. **(B)** Energy versus net charge for the same random ensemble. **(C)** Total-energy distribution of 10,000 random decoys compared with 1,000 locally optimized sequences; the optimized basin is shifted toward lower Z-scores. **(D)** Average helical-wheel representation of the optimized basin, showing ensemble-level amphipathic segregation. **(E)** Hydrophobicity autocorrelation across the optimized sequences, with peaks at *k* = 3 and *k* = 4 consistent with *α*-helical periodicity.

## 2 Discussion

This work introduces a physics-based Hamiltonian framework for environment-conditioned design of membrane-parallel *α*-helical peptides. The central contribution is to make the environment part of the sequence objective itself. Helix propensities, local sequence context, pairwise contacts, solvent polarity, membrane charge coupling and electrostatic screening are represented as explicit terms, and their contributions are placed on a common statistical scale through random-reference Z-score normalization. The resulting score is therefore both interpretable and portable across sequence lengths, amino-acid alphabets and physical environments.

This formulation occupies a distinct position within the current biomolecular modeling landscape. Rosetta-like protocols, atomistic simulations and deep-learning methods have transformed structure prediction, scaffold generation, sequence design and detailed refinement [23, 9, 10, 24, 25]. The present model addresses a more specific computational problem: optimization of sequence identity on a prescribed helical scaffold under controllable environmental constraints. In this setting, solvent polarity, surface charge, amphipathic exposure and membrane-facing geometry can be varied directly in the objective. This is important for membrane-active peptides, where the relevant design criterion is not generic helicity, but relative compatibility with one environment over competing aqueous, apolar or interfacial alternatives.

The benchmarks support this interpretation at three levels. First, experimentally characterized water-soluble *α*-helical peptides and WALP/GWALP transmembrane model peptides are assigned the expected homogeneous-medium preference. Second, canonical membrane-active amphipathic peptides are recognized as favorable at anionic interfaces, whereas non-AMP helix-forming controls remain more compatible with water. Third, prospective objectives generate distinct sequence families for cross-environment compatibility and anionic one-vs-rest selectivity. Together, these tests show that favorable scores reflect environment-specific organization rather than helix propensity alone.

The landscape analyses provide an additional check on the design behavior. Low-energy sequences show increased transverse hydrophobic moments, controlled charge distributions, a separated optimized basin and *i, i*+ 3/*i*+ 4 hydrophobicity periodicity, even though hydrophobic moment is not included as an explicit energetic term. These emergent features indicate that the combined Hamiltonian terms are sufficient to organize the sequence landscape around chemically interpretable amphipathic patterns.

A practical advantage of the formulation is its direct connection to combinatorial optimization. Because the model is written as a discrete Hamiltonian over amino-acid identity variables, the same physical objective can be explored by exact enumeration for small systems, classical heuristics for larger searches, or hybrid quantum-classical algorithms after binary encoding. This quantum-ready structure does not imply a demonstrated quantum speedup in the present work, but it provides a clear route for implementing the same environment-conditioned design objective on qubit-based optimization backends without changing the biophysical model.

The present scope is sequence-level, environment-conditioned scoring on a prescribed *α*-helical scaffold. For interfacial cases, penetration depth and orientation are treated as explicit geometric variables: a sequence can be evaluated across this grid, and its preferred interfacial placement is identified from the corresponding Z-score landscape. The generated candidates should therefore be treated as computational priorities for follow-up validation rather than experimentally established leads. Future work should combine this Hamiltonian with stronger compositional constraints, atomistic or coarse-grained structural validation and direct experimental tests. In this role, the framework provides a transparent computational-science benchmark for environment-conditioned peptide design and a practical route to selective candidate generation.

## 3 Methods

A binary-encoded Hamiltonian framework is used for the design of membrane-interacting *α*-helical peptide sequences on a fixed scaffold. The formulation encodes peptide sequences using a compact binary representation, where each position *i* (for *i* = 1, …, *L*) is assigned b = ⌈log_2_|𝒜|⌉ bits to represent one of |𝒜| amino acids in the alphabet 𝒜. Projector operators enforce selection of valid amino acid codes, effectively mimicking one-hot encoding through products of (1 ± Z_*k*_)/2 terms in the qubit basis, where Z_*k*_ are Pauli-Z operators acting as projectors onto the |0⟩ or |1⟩ eigenstates on qubit k [26, 27, 28]. For conceptual clarity, the Hamiltonian is described in terms of effective one-hot variables *x*_*i,α*_ ∈ {0, 1}, where *x*_*i,α*_ = 1 if amino acid *α* is selected at position *i*, with the constraint Σ*α x*_*i,α*_ = 1 enforced via penalties.

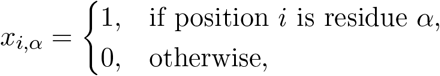

where *i* = 1, …, *L* and *α* ∈ 𝒜.

The total Hamiltonian is decomposed into multiple components capturing residue-level helix propensities, local sequence-context effects, pairwise interactions, membrane-specific effects, and structural constraints:

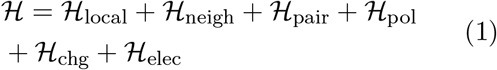

### 3.1 Hamiltonian Formulation for Environment-Conditioned Helical Design

#### Local contributions ℌ_local_

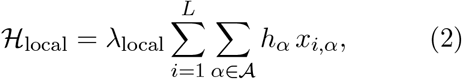

In this expression, *h*_*α*_ represents a fixed energetic contribution assigned to each amino acid type *α*, capturing its intrinsic *helix-forming propensity*. In this work, these values are taken directly from the empirical helix-propensity scale reported by Pace and Scholtz [29], which quantifies the free-energy changes associated with helix formation.

This term energetically favors residues with high helical stability (e.g., Alanine or Glutamic acid) uniformly across all sequence positions. Representative propensity values used in this model are detailed in Supplementary Table S1. Effectively, ℌ_local_ serves as a residue-specific background energy term, a common feature in statistical and physical protein-energy Hamiltonians used for fold recognition and design.

#### Helix Neighbor Interaction Terms (ℌ_neigh_)

To capture how local sequence context modulates helical stability beyond individual residue propensities, short-range neighbor interactions based on empirical statistical matrices are included [30]:

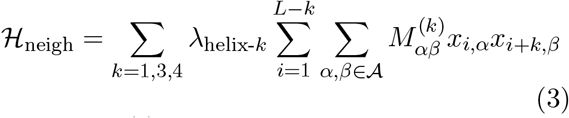

where *M*^(*k*)^ represents the normalized interaction matrices for the first (*k* = 1), third (*k* = 3), and fourth (*k* = 4) neighbors [31]. These matrices reflect the statistical preference for residue pairs at specific sequence separations to stabilize or destabilize the helical fold.

Notably, the *k* = 2 interaction is omitted from this summation. Due to the periodicity of the *α*-helix (~3.6 residues per turn), residues at positions *i* and *i* + 2 are oriented on nearly opposite faces of the helix (approximately 200*°* apart). Consequently, their side chains are physically sequestered from one another, making significant direct interactions negligible compared to the proximal *i* + 1 pairs or the face-aligned *i* + 3 and *i* + 4 pairs[32].

#### Pairwise interactions ℌ_pair_

Pairwise terms of the form:

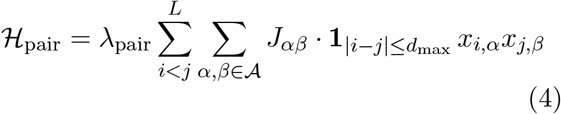

Where the components of the equation are defined as follows:

- λ_**pair**_: A global scaling factor that modulates the relative weight of pairwise interactions within the total Hamiltonian.
- *J*_*αβ*_: Represents the interaction energy between residue types *α* and β. These values are drawn from the Miyazawa-Jernigan contact matrix [33]. Such effective pair potentials are foundational in protein modeling and remain central to fold recognition and design force fields [33, 34].
- **1**_|*i−j*|*≤d*max_ : This is the indicator function (or characteristic function). It returns 1 if the sequence separation between residues at positions *i* and *j* is within the threshold *d*_max_ (default value of 4, corresponding to pairs from *i, i* + 1 through *i, i* + 4 along the peptide chain), and 0 otherwise.

#### Environment-Mediated Interactions (ℌ_pol_ and ℌ_chg_)

To account for the anisotropic nature of the membrane interface, the environmental energy is decomposed into two complementary terms: a solvation-based polarity term, ℌ_pol_, and an electrostatic coupling term, ℌ_chg_. These terms collectively drive the formation of amphipathic structures by partitioning residues according to the helical phase.

#### 1. Solvation Polarity and Amphipathicity (ℌ_pol_)

This term models the energetic cost of burying polar residues or exposing hydrophobic ones relative to the membrane-water interface. The energy is defined as:

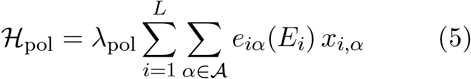

The local environment of residue *i* is determined by its angular position on the helical wheel:

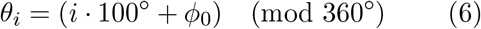

where 100*°* is the angular advance per residue in an ideal *α*-helix, corresponding to approximately 3.6 residues per turn, and ϕ_0_ is the helical-wheel phase. Membrane exposure is parameterized by an external penetration variable *p* through the membrane-facing angular half-width

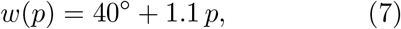

with *p* expressed in percent. A discrete exposure state *E*_*i*_ ∈ {membrane, water} is then defined for each residue: residues satisfying |*θ*_*i*_| ≤ *w*(*p*) are assigned to the membrane phase, and the remaining residues are assigned to the water phase. Larger values of *p* therefore broaden the non-polar exposure sector. In a given calculation, *p* is treated as an external geometric parameter that can either be fixed to define a design objective or scanned to evaluate a known sequence; the optimizer acts only on sequence identity and does not vary insertion depth, helix orientation, or the exposure pattern.

For **membrane-facing** positions (*E*_*i*_ = membrane):

- If *α* is polar, *e*_*iα*_ assigns a desolvation penalty (+|*H*_*α*_|).
- If *α* is hydrophobic (*H*_*α*_ > 0), *e*_*iα*_ provides a favorable partition reward (−|*H*_*α*_|).

For **water-facing** positions (*E*_*i*_ = water), these preferences are inverted. Hydrophobicity values *H*_*α*_ are drawn from the Fauchère-Pliska scale [35].

#### 2. Membrane Charge Surface Coupling (ℌ_chg_)

Complementary to the hydrophobic effect, the model incorporates the electrostatic attraction or repulsion between amino acid side chains and the net charge of the lipid headgroups:

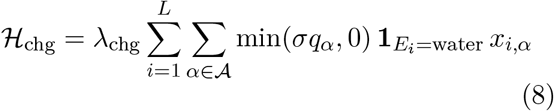

where *q*_*α*_ is the formal charge of residue *α* and σ ∈ {− 1, 0, 1} represents the membrane surface potential (negative, neutral, or positive). The indicator function 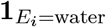 restricts this interaction to residues oriented towards the aqueous interface where external ionic coupling occurs. The min(σq_*α*_, 0) factor makes this term explicitly unidirectional: only favorable opposite-sign couplings contribute, whereas same-sign couplings contribute zero rather than a positive penalty. This coarse-grained choice captures selective charge complementarity of amphipathic peptides at charged interfaces without introducing an additional repulsive surface term. In this work, Lys and Arg are assigned *q* = +1, Asp and Glu are assigned *q* = −1, and histidine is treated as neutral in the charge-dependent terms.

#### Electrostatic Screening Terms ℌ_elec_

Electrostatic interactions between residue charges are modeled using an effective sequence-separation- and environment-dependent screening factor, consistent with coarse-grained and implicit-solvent approaches [36, 37]. The electrostatic contribution is

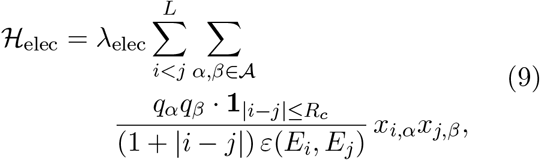

where *q*_*α*_ and *q*_*β*_ are the charges of residue types *α* and β, and *x*_*i,α*_ indicates residue identity. The cutoff *R*_*c*_ = 8 is measured in residue-index steps along the peptide chain; it limits the electrostatic term to pairs from *i, i*+1 through *i, i*+8, while the interaction strength within this window decays as 1/(1 + | *i* − *j* |) [38, 39]. The notation *R*_*c*_ is used rather than *d*_max_ to distinguish this longer electrostatic screening range from the shorter generic contact window in ℌ_pair_. The effective dielectric constant ε(*E*_*i*_, *E*_*j*_) explicitly depends on the discrete exposure states of both interacting residues: ε = 0.1 when both residues are in the membrane phase (*E*_*i*_ = *E*_*j*_ = membrane), ε = 1.0 when both are in the solvent (*E*_*i*_ = *E*_*j*_ = water), and ε = 0.3 for mixed interfacial pairs, following established coarse-grained membrane parameters [37, 36].

### 3.2 Transverse Hydrophobic-Moment Descriptor

Helix amphiphilicity is quantified through the transverse hydrophobic moment of Eisenberg *et al*. [40]. This descriptor is used as a reaction coordinate in the landscape analysis and as an orientation rule when aligning benchmark peptide sequences before the interfacial penetration scan.

Let 𝒜 be the amino-acid alphabet and *L* the helix length. The projected components of the hydrophobic moment are defined as:

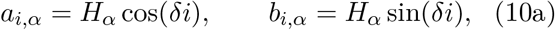

where *H*_*α*_ is the hydrophobicity value (see Supplementary Table S2), and δ = 100*°* for an *α*-helix. Summing yields

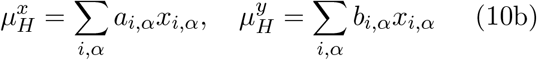

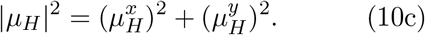

The magnitude |µ_*H*_| is evaluated after sequence generation as an amphipathicity observable, and its direction is used in the membrane-active peptide benchmark described below to define the membrane-facing orientation before scanning penetration.

### 3.3 Thermodynamic Consistency and Statistical Normalization

A critical challenge in constructing a multi-term Hamiltonian for protein design is the heterogeneity of energy scales and their dependence on the sequence length *L*. Without correction, the optimization landscape suffers from two primary artifacts: *Thermodynamic Inconsistency*, where the relative weight of local vs. pairwise terms shifts with *L*, and *Gradient Bias*, where terms with larger numerical magnitudes dominate the optimization process [41].

To resolve these issues, a two-stage normalization protocol is implemented based on intensive energy descriptors and Z-score transformations in the context of spin-glass theory [18].

#### 1. Length-Dependent Scaling

To keep each energy contribution intensive with respect to sequence length *L*, specific normalization factors are applied to each class of interaction. This prevents the over-representation of long-range or high-frequency terms in larger systems and mitigates the risk of scale dominance [41]. The following scaling factors are applied:

- **Local terms (1-body):** Standardized by the total sequence length:

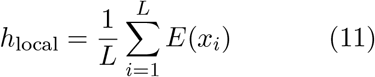
- **Restricted pairwise interactions:** Scaled by the number of active pairs at a fixed sequence separation *k*:

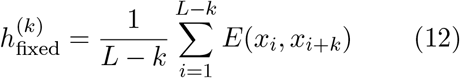
- **Neighboring contact pairs:** For interactions within a defined range *k*_max_ (e.g., sequence neighbors *i, i* + 1, …, *i* + k):

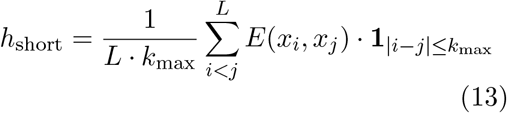
- **Global (All-to-All) interactions:** Scaled by the total number of possible unique pairs 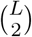 to maintain a consistent average energy per pair:

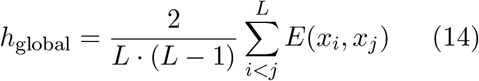

By applying these specific denominators, the average energy density of each physical effect is preserved regardless of peptide size, making the scaled objective comparable across design targets and optimization backends[41].

#### 2. Z-Score Transformation via Decoy Sets

Second, to eliminate Gradient Bias and establish a physically meaningful reference state, each Hamiltonian term_*k*_ is transformed into a dimensionless Z-score space. For each term, the mean µ_*k*_ and standard deviation σ_*k*_ are estimated by evaluating a random decoy set of *N*_decoy_ non-designed sequences, with typical values ranging from approximately 10,000 to 60,000 sequences depending on the analysis. The sampling size is reported explicitly in each benchmark and can be configured in the web implementation:

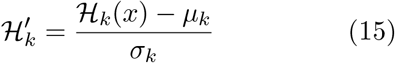

where 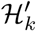 represents the energy in units of standard deviation from the random background. This transformation provides three crucial advantages:

- **Iso-Variance (**σ = 1): All energy terms are normalized to unit variance, ensuring an equal “statistical force” during optimization.
- **Reference State (ℌ***′* ≈ 0): A zero value represents the energy of the random-reference ensemble.
- **Stability Criterion:** Sequences with ℌ*′* < 0 are identified as energetically favorable within the model, while ℌ*′* > 0 represents unfavorable, high-energy configurations.

Unless otherwise noted, all Z-scores reported below are therefore expressed in units of the corresponding random-ensemble standard deviation σ. The corresponding raw and standardized term distributions are provided in the Supplementary Information. Before standardization, the raw Hamiltonian terms span markedly different numerical ranges, so terms with larger baseline variance dominate the objective. After Z-score normalization, each contribution is placed on a common dimensionless scale relative to a random decoy background, making the resulting multi-term Hamiltonian easier to interpret and balance across sequence lengths.

The final multi-objective Hamiltonian is then constructed as the weighted sum of these standardized terms: 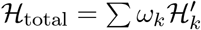. This formulation allows the algorithm to navigate a balanced landscape where convergence is driven by physical stability rather than numerical scale[18, 42, 43].

### 3.4 Weighting Scheme and Model Parameterization

The normalization procedure above places all Hamiltonian components on a common statistical scale, but it does not by itself determine their final relative importance. The weights λ_*i*_ therefore play a distinct methodological role: they define how strongly the normalized physical effects compete once raw magnitude differences and length dependence have been removed. In this work, these weights were not fitted to maximize performance on any single downstream benchmark. Instead, they were chosen to preserve a physically balanced competition among solvation, charge, electrostatic, local-propensity, pairwise, and neighbor-dependent terms after normalization, while avoiding trivial domination by any one contribution.

For the full Hamiltonian used in the main analyses, the explicit effective weights listed in the Supplementary Information are used. This parametrization was stabilized empirically by requiring three conditions simultaneously: first, the normalized terms should remain of comparable order across random-sequence backgrounds; second, the resulting Hamiltonian should recover the expected directional behavior in homogeneous polar and apolar limits; and third, interfacial designs should display chemically sensible amphipathic segregation without requiring any explicit hydrophobic-moment energy term.

The resulting hierarchy is physically interpretable. The polarity term receives the largest weight (λ_pol_ = 6) because solvent partitioning is the dominant driver of amphipathic segregation at an interface. The electrostatic pair term remains strong (λ_elec_ = 4) because it suppresses unrealistically overcharged designs and favors alternating charge patterns in the aqueous sector, while the membrane-surface charge coupling is somewhat smaller (λ_chg_ = 3) because it acts only on water-facing residues and only through favorable sign complementarity. By contrast, the local helix-propensity and neighbor terms are assigned moderate weights (λ_local_ = λ_neigh_ = 1.5): they are needed to bias the search toward coherent helical sequences, but they should not overwhelm the environmental physics. Finally, the generic pairwise contact term is kept weak (λ_pair_ = 0.1), so that it serves mainly as a soft compositional regularizer rather than as the primary determinant of amphipathic patterning. The final values should therefore be interpreted as a physically ranked parametrization, not as a benchmark-specific fit.

The adequacy of this weighting scheme is then assessed a posteriori in the Results section through several independent tests: recovery of the homogeneous polar and apolar limits, recognition of known water-soluble and transmembrane model peptides in the corresponding homogeneous media, recognition of canonical membrane-active peptides at anionic interfaces, and the emergence of both cross-environment and one-vs-rest selective design regimes. In that sense, the weights are justified methodologically by balance and physical ranking, and validated subsequently by the coherence of the resulting design behavior across independent tasks.

### 3.5 Binary Encoding and Algorithmic Portability

The normalized Hamiltonian defined above extends beyond a simple cost function for deterministic optimization; it is mathematically isomorphic to a generalized Potts model [17]. In statistical mechanics, such formalisms define an energy landscape where the probability of observing a specific microstate (a peptide sequence *S*) at thermal equilibrium is governed by the Boltzmann distribution:

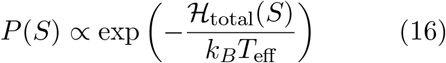

where *k*_*B*_*T*_eff_ represents an effective computational temperature regulating the exploration-exploitation trade-off.

Unlike evolutionary statistical models—such as Direct Coupling Analysis (DCA) [44] or EVmutation [45]—which infer pairwise couplings (*J*_*ij*_) and local fields (*h*_*i*_) inversely from multiple sequence alignments, the present Hamiltonian formulation constitutes a *forward* predictive physical model. The interactions are explicitly derived from biophysical principles instead of historical evolutionary data.

Consequently, executing the optimization algorithm (whether via classical Simulated Annealing or quantum QAOA) does not merely return a single, functionally isolated global minimum. Instead, the solvers act as statistical samplers, generating an *ensemble* of probable sequence candidates residing in the low-energy basins of the landscape. The Z-score standardization serves as an empirical normalization over the unfolded reference state, ensuring that the generated ensembles correspond to thermodynamically favorable configurations within the model. On that basis, the model is implemented through a compact binary encoding that preserves the physical Hamiltonian while improving algorithmic portability across classical and quantum-compatible backends.

In sequence-design QUBO formulations, one-hot encoding enforces that each sequence position hosts exactly one amino acid. This constraint is typically implemented by adding a quadratic penalty of the form (1 – Σ_*α*_ *x*_*i,α*_)^2^ per site. Such encodings are standard in combinatorial optimization and widely used in lattice protein models; e.g., Irbäck *et al*. mapped the HP-model design problem to an Ising spin glass with a one-hot choice at each site [46].

#### 3.5.1. Binary Encoding Implementation

To mitigate the scalability limitations inherent in traditional one-hot encoding, which requires 𝒪 (*L*|𝒜|) qubits, a binary encoding scheme is implemented. This strategy significantly optimizes quantum resource allocation by reducing the qubit overhead to 𝒪 (*L* log_2_ |𝒜 |), where L denotes the sequence length and represents the amino acid alphabet, defined as:

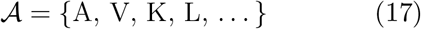

This logarithmic scaling is essential for mapping protein design problems onto near-term quantum hardware with limited qubit counts [21].

#### Qubit Allocation

Each position *i* in the protein sequence uses *b* = log_2_|𝒜| qubits to encode the amino acid selection. The global qubit index for bit *k* at position *i* is calculated as:

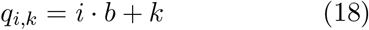

For a sequence of length *L*, the total number of qubits required is:

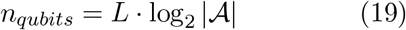

#### Amino Acid Mapping

Each amino acid *α* ∈ {0, 1, …, |𝒜| −1} is mapped to its binary representation. For example, with 4 amino acids (|𝒜| = 4, *b* = 2):

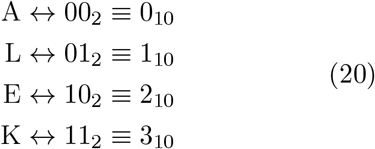

To facilitate a rational selection of amino acids based on their local properties, the amino-acid set was analyzed in a normalized physicochemical space using both Principal Component Analysis (PCA) and pairwise distance diagnostics. These supplementary similarity maps provide an interpretable view of biochemical redundancy and motivate the reduced alphabets used in the larger calculations; the corresponding plots are provided in the Supplementary Information.

#### Dimensionality Reduction via *k*-means Clustering

To systematically reduce the dictionary size |𝒜| for hardware compatibility without losing chemical diversity, a *k*-means clustering algorithm was applied to the normalized 3D property space (hydrophobicity, helix propensity, and charge). By defining the number of clusters *k* (e.g., *k* = 4, 5, or 8), the continuous space is partitioned into *k* distinct physicochemical regions.

To ensure that the quantum algorithm evaluates real molecular entities, the representative amino acid for each cluster was selected by finding the residue with the minimum Euclidean distance to the computed cluster centroid. Proline was manually excluded from the reduced dictionaries due to its unique helix-breaking properties. This dimensionality reduction ensures the problem remains computationally tractable under the 𝒪 (*L* log2|𝒜|) binary encoding scheme while preserving the essential biophysical features required for stable folding.

#### 3.5.2 Hamiltonian Construction via Projector Expansion

The binary encoding necessitates expressing amino acid preferences through projector operators. For amino acid *α* at position *i* with preference coefficient *h*_*α*_, the projector 𝒫_*α,i*_ selects the corresponding binary state.

##### Single-Qubit Projectors

For each bit *k* with value *v*_*k*_ ∈ {0, 1} in the binary representation of *α*, the projector onto eigenvalue 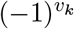 of Pauli-Z is:

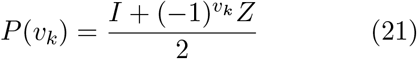

##### Multi-Qubit Projector

The full projector for amino acid *α* at position *i* is:

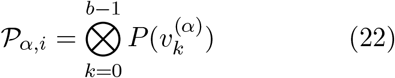

where 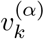 is the *k*-th bit of *α*’s binary representation.

##### Pauli Expansion

Expanding the projector yields 2_*b*_ terms:

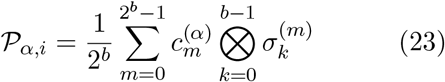

where 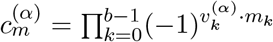 and 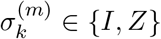 depending on bit *m*_*k*_ of mask *m*.

It is critical to note that while traditional one-hot encodings yield a strict Quadratic Unconstrained Binary Optimization (QUBO) problem with at most two-body interactions, the compact binary encoding theoretically introduces higher-order terms (multi-qubit interactions up to order 2*b*) upon projector expansion. These higher-order terms are natively and efficiently explored by gate-based quantum algorithms such as QAOA. However, for direct execution on classical or quantum annealing hardware (e.g., D-Wave), which natively accept only quadratic topologies, the resulting Hamiltonian must either be quadratized using ancilla variables or executed in the standard one-hot representation.

#### 3.5.3 Solution Decoding

Quantum measurement outcomes are decoded by converting binary qubit states to amino acid indices:

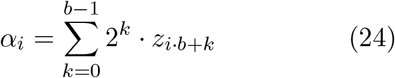

where *z*_*j*_ ∈ {0, 1} represents the measured state of qubit *j*.

#### 3.5.4 Scalability Analysis

The binary encoding achieves significant qubit reduction compared to one-hot encoding:

To quantify the scaling of the search space, peptide sequences of length *L* = 10 were considered using a reduced amino acid alphabet of size |𝒜| = 16. The total number of distinct sequences is

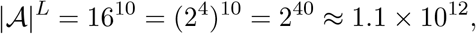

corresponding to approximately 1.1 trillion unique candidate sequences. Because 16 residue types require exactly 4 bits per position, each chemically valid sequence maps one-to-one onto a 40-bit assignment. Thus, an exhaustive search over the reduced alphabet and an exhaustive search over the corresponding binary strings involve the same number of candidates.

Even under an optimistic order-of-magnitude assumption in which the evaluation of the full Hamiltonian energy for a single sequence requires only τ ≈ 10^*−*4^ seconds (0.1 ms), the total wallclock time required for an exhaustive brute-force enumeration would be

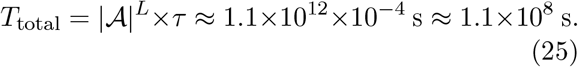

This corresponds to approximately 3.5 years of uninterrupted single-core computation for a single design instance. This estimate is intended only as an illustrative order-of-magnitude benchmark for the exponential bottleneck, not as a hardware-specific runtime claim for a particular implementation. This exponential scaling illustrates a fundamental computational bottleneck: for sequence lengths *L* ≥ 10, exact classical enumeration rapidly becomes impractical. Consequently, practical peptide design at this scale generally requires stochastic sampling strategies or heuristic approaches, including the hybrid quantum–classical framework employed in this work.

Parallelization mitigates this cost only linearly and does not alter the underlying exponential scaling with *L*[47, 48].

For compact binary-encoded optimization runs, a 16-amino-acid working alphabet requiring 4 qubits per residue was used. This set was obtained by merging near-redundant amino-acid types in the underlying physicochemical space, yielding { *A, P, D, E, Q, L, M, F, K, S, T, W, Y, V, H, G*}.

This reduction is a computational choice for selected optimization tasks, while the scoring model also supports larger alphabets. Other analyses in the manuscript, including reference-sequence scoring and membrane-active peptide benchmarks, use larger alphabets up to the full 20 amino acids and estimate random-reference Z-scores from decoy samples. The web implementation follows the same principle for user-specified sequences and alphabets, with configurable decoy sampling. For longer-sequence exploratory calculations discussed later and in the Supplementary Information, additional *k*-means reductions to smaller dictionaries (e.g., *k* = 4 or *k* = 8) were also used to further compress the search space.

### 3.6 Quantum and Classical Optimization Backends

Three optimization backends were employed to explore the sequence landscape defined by the Hamiltonian: exact or heuristic classical search, simulated annealing with D-Wave Neal, and the Quantum Approximate Optimization Algorithm (QAOA). These methods serve different purposes within the study. Classical enumeration provides ground truth for small systems, simulated annealing extends the exploration to larger instances, and QAOA tests whether the same physically motivated objective can be handled within a hybrid quantum-classical workflow. The implementation uses Qiskit v1.4.5[49] as the primary development kit, with an optional PennyLane backend[50]. Post-processing of measurement outcomes, probability distributions, and sequence decoding was performed with Python tools based on NumPy [51] and Matplotlib[52].

#### 3.6.1 Quantum Approximate Optimization Algorithm (QAOA)

QAOA was used as a hybrid quantum-classical backend for optimizing the same binary-encoded Hamiltonian explored by the classical solvers[53, 54]. A schematic overview of the workflow is provided in the Supplementary Information.

In this implementation, the ansatz consists of *p* layers, each applying a unitary evolution driven by the cost Hamiltonian followed by a mixing Hamiltonian. The cost Hamiltonian encodes the protein folding energy landscape, while the mixer introduces superposition to explore the computational basis.

For small systems (*L* = 2), a linear ramp initialization strategy[55, 56] was employed to guide the optimization towards the ground state, setting the circuit depth to *p* = 5. For larger systems (*L* ≥ 8), a standard random initialization with *p* ∈ [1, 2] layers was used. The variational parameters are optimized using COBYLA, a gradient-free optimizer[57]. In this regime, COBYLA performs a stochastic local search, greedily minimizing energy from an arbitrary starting point. To mitigate the risk of becoming trapped in suboptimal local minima, a multi-start strategy is used, executing independent optimization runs to sample diverse regions of the solution space.

A Conditional Value at Risk (CVaR) objective function is used to prioritize the lower tail of the measured energy distribution, significantly improving the probability of finding the ground state sequence compared to standard expectation value minimization[58].

#### 3.6.2 Simulated Annealing Heuristic

`dwave-neal` was used as a classical simulated annealing (SA) implementation for heuristic exploration of larger binary design instances. neal operates on classical CPUs, performing local spin-flip updates guided by the Metropolis acceptance criterion. Unlike statevector-based quantum simulations, which scale exponentially as *O*(2^*N*^)[59], Neal scales linearly in memory *O*(*N*)[60], allowing efficient exploration of peptide-design instances beyond the limits of exact quantum simulation. This provides a practical heuristic baseline for evaluating the energetic consistency of the Hamiltonian[60, 61, 62].

#### 3.6.3 Classical Benchmark Solver

To validate the quantum results, a classical solver was implemented that treats the protein design problem as a Combinatorial Optimization task. The Hamiltonian is evaluated purely as a classical cost function 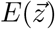, where 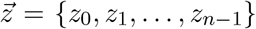 represents the vector of binary variables defined in the Binary Encoding section.

For tractable problem sizes (up to *n* = 20 binary variables), the solver performs an exhaustive brute-force enumeration of the entire solution space (2^*n*^). This approach guarantees the identification of the global minimum bitstring 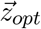 and provides an exact reference spectrum for evaluating the other optimization backends.

For larger instances where the search space size (2^*n*^) renders exhaustive enumeration intractable, the solver utilizes a heuristic strategy combining Massive Random Sampling followed by a Hill Climbing local search algorithm[63]. This two-stage approach efficiently explores the energy landscape to find high-quality local minima, serving as a robust baseline to assess the performance and accuracy of the quantum solvers in complex regimes.[64].

#### 3.6.4 Role of Shots

In both quantum solvers, shots correspond to the number of times a circuit is executed to obtain measurement statistics. Because quantum measurements are inherently probabilistic, multiple shots are necessary to accurately estimate expectation values of the Hamiltonian and marginal probabilities. These statistics provide both the estimated ground-state energy and a probability distribution over candidate sequences, facilitating the selection of feasible protein sequences with high likelihood.

### 3.7 Benchmark and Design Protocols

#### 3.7.1 Benchmarking Logic and Negative Controls

The validation strategy was designed to separate four questions that are often conflated in peptide-design workflows. First, random-reference Z-scores test whether a sequence is unusually favorable relative to unconstrained sequence backgrounds of the same length and alphabet. Second, reference-helix recognition tests whether the Hamiltonian assigns the expected environmental preference to experimentally characterized sequence classes that were not generated by the optimizer. Third, negative-control sequences test whether favorable interfacial scores reflect membrane-active amphipathic organization rather than generic helix propensity. Fourth, landscape diagnostics test whether the optimized basin has coherent collective structure, including amphipathic segregation and helix-periodic hydrophobicity patterns, even though the hydrophobic moment is used only as a diagnostic or alignment descriptor and not as an explicit energy term.

#### 3.7.2 Aqueous and Apolar-Core Reference-Helix Benchmark

To benchmark homogeneous polar/apolar scoring without treating the optimized boundary cases as experimentally validated structures, reference peptides whose environment compatibility is known a priori were evaluated. The aqueous set comprised alanine-rich Glu/Lys peptides from the Marqusee–Baldwin and Scholtz– Baldwin families of short water-soluble helix studies[65, 66, 67]. The apolar-core set comprised WALP and GWALP transmembrane model peptides, including Tyr-anchor GWALP variants, which are established hydrophobic *α*-helical peptides in lipid-bilayer environments[68, 69, 70]. Their helical assignments come from the original experimental characterization: circular-dichroism and helix–coil analyses for the alanine-rich aqueous peptides, and circular-dichroism together with solid-state NMR/oriented-bilayer measurements for the WALP/GWALP membrane peptides. Here these transmembrane model peptides are used as apolar-core references, not as literal homogeneous-solvent standards.

Each sequence was treated as a fixed helical primary structure rather than as a design variable, and terminal modifications reported for some reference peptides were not included in the sequence-only Hamiltonian. The same amino-acid sequence was scored in the homogeneous polar and homogeneous apolar presets using fixed environments, with final Z-scores computed against 10,000 random sequences of matching length and alphabet. The reported quantity is Δ*Z* = *Z*_apolar_ − *Z*_water_, so that positive ΔZ indicates water recognition and negative ΔZ indicates apolar-core recognition. This benchmark tests whether the Hamiltonian recognizes known environment-compatible helix classes; it is not a de novo folding or atomistic stability calculation.

#### 3.7.3 Interfacial Benchmark Protocol

For the membrane-recognition benchmark, seven canonical membrane-active amphipathic peptides (`LL-37`, `Melittin`, `Magainin 2`, `Cecropin A`, `PGLa`, `Piscidin 1`, and `BP100`) were evaluated together with two non-AMP helix-forming controls (`(EAAAK)3` and `Poly-Ala 15`). Each sequence was treated as a fixed *α*-helical peptide of known primary structure rather than as a design variable. Before scanning penetration, the helical-wheel phase was aligned so that the transverse hydrophobic moment points toward the non-polar side of the interface. This alignment uses µ_*H*_ only as a geometric orientation rule and not as an energetic contribution.

For each aligned sequence, the anionic, neutral, and cationic interfacial Hamiltonians were then evaluated over sampled penetrations from 5% to 95% in 5% increments, corresponding to exposure masks generated through w(*p*) = 40*°* + 1.1 p. The reported quantities *Z*_water_ and *Z*_apolar_ correspond to the homogeneous polar and homogeneous apolar Hamiltonians, whereas 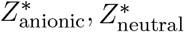, and 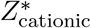 denote the most favorable interfacial Z-scores obtained over the penetration scan. The associated *p*^*∗*^ values identify the sampled penetration at which each interfacial optimum is found. Unless otherwise stated, these benchmark Z-scores are referenced to 3,000 random sequences of the same length over the full 20-amino-acid alphabet.

#### 3.7.4 Multi-Environment Design Objectives

To probe multi-environment robustness, sequences were optimized against the joint homogeneous objective

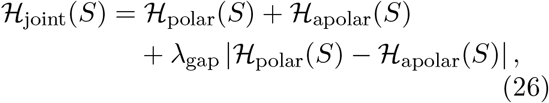

where λ_gap_ penalizes imbalance between the two media. The cross-environment calculations reported in the main text use *L* = 20, the reduced 16-residue working alphabet, and a neutral treatment of histidine in the charge-dependent terms. Final sequence energies are converted to homogeneous-medium Z-scores relative to random decoys generated with the same length and alphabet.

For explicit one-vs-rest selectivity, a target environment 𝒯 and a competing set of off-target environments 𝒪 are defined. In the present manuscript, the proof-of-principle target is an anionic membrane interface, reflecting the negatively charged membrane surfaces commonly associated with bacterial and other diseased-cell targets of membrane-active peptides [4, 5, 8]. The off-target set is {neutral interface, cationic interface, homogeneous water, homogeneous apolar medium}, representing competing environments that should not be preferentially stabilized by an anionic-membrane-selective design. Because interfacial energetics depend on the exposure mask, these selectivity calculations are carried out at fixed imposed penetrations rather than after a post hoc scan. The interpolating objective family is therefore

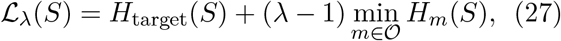

with 0 ≤ λ < 1. This compact form is algebraically equivalent to λH_target_(*S*) + (1 − λ) [*H*_target_(*S*) − min_*m∈𝒪*_ *H*_*m*_(*S*)], but makes the target-versus-margin interpolation more transparent. For the present single-target case this becomes 𝔏_*λ*_(*S*) = *H*_anionic_(*S*) + (λ −1min(H_neutral_, *H*_cationic_, *H*_polar_, *H*_apolar_). Thus λ = 0 recovers the hard max–min selectivity limit, whereas larger λ values progressively place more emphasis on absolute target favorability relative to the one-vs-rest margin. In the current study, this objective is evaluated for fixed anionic penetrations of 10%, 25%, 50%, 75%, and 90% using *L* = 20 sequences over the full 20-amino-acid alphabet and the values λ = 0, 0.25, 0.5, and 0.75.

## 4 Availability of Data and Software

https://simbios.usc.es/HelixDesignStudio/

## 5 Declaration of AI and AI-Assisted Technologies in the Writing Process

During the preparation of this work, the authors used ChatGPT (OpenAI, GPT-5) to improve language and readability. After using this tool, the authors reviewed and edited the content as needed and take full responsibility for the content of the publication.

## Acknowledgments

This work was supported by the Spanish Agencia Estatal de Investigación (AEI) and the ERDF (PID2022-141534OB-I00 and CNS2023-144353), by Xunta de Galicia (ED431C 2025/15, ED431C 2021/21 and Centro de investigación do Sistema universitario de Galicia accreditation 2023–2027, ED431G 2023/03) and the European Union (European Regional Development Fund – ERDF). Authors also acknowledge funding from RePoSUDOE, with project reference S1/1.1/P0033, a project co-financed by the Interreg Sudoe Programme through the European Regional Development Fund (ERDF). D.C.T. thanks the Ministerio de Universidades for his predoctoral contract (FPU22/00636). Computational calculations in this work were performed at CESGA.

## Notes

https://simbios.usc.es/HelixDesignStudio/

### Competing Interest Statement

The authors have declared no competing interest.

