## Supplementary Information for "Environment-conditioned design of *α*-helical peptides"

This Supplementary Information provides implementation details and extended results supporting the computational benchmark hierarchy described in the main text. Sections S1–S3 summarize the Hamiltonian terms, normalization strategy, benchmark structure, and the supplementary evidence linked to each main-text claim. Sections S4–S10 document the quantum-compatible optimization backend, parameter initialization, classical variational optimization, reduced-alphabet construction, and solution-selection workflow. Sections S11–S14 report empirical amino-acid parameters, Hamiltonian normalization and weighting details, supporting methodological figures, boundary-case sanity checks, additional low-energy landscape diagnostics, reference-sequence recognition tables, and complete one-vs-rest anionic-specificity design libraries. The supplementary results support reproducibility and interpretation of the Hamiltonian benchmark; generated sequences remain computational predictions until tested experimentally or by independent structural simulation.

#### S1. Hamiltonian Components and Physical Roles

The main text defines a single sequence Hamiltonian reused across homogeneous aqueous, homogeneous apolar, neutral interfacial, cationic interfacial, and anionic interfacial environments. The model combines six energetic contributions: residue-level helical propensity, neighbor-dependent helical context, local pairwise contacts, solvent-polarity preference, environment-dependent charge coupling, and screened electrostatics. The transverse hydrophobic moment is reported as an amphipathicity descriptor and alignment coordinate in selected benchmark scans.

Rebeca García-Fandiño:

Ángel Piñeiro:

Table S1: **Hamiltonian terms used across the main analyses.** Each term is standardized against random sequence backgrounds before weighted combination.

| Term | Physical role |
|---|---|
| $\mathcal{H}_{\text{local}}$ | Residue-specific $\alpha$-helical propensity on the fixed helical scaffold. |
| $\mathcal{H}_{\text{neigh}}$ | Neighbor-dependent helical compatibility for local sequence contexts. |
| $\mathcal{H}_{\text{pair}}$ | Short-range residue-pair compatibility within local sequence neighborhoods. |
| $\mathcal{H}_{\text{pol}}$ | Compatibility with polar, apolar, or interfacial exposure states. |
| $\mathcal{H}_{\text{chg}}$ | Coupling between residue charge and the specified membrane or solvent environment. |
| $\mathcal{H}_{\text{elec}}$ | Screened electrostatic interaction between charged residues under the local exposure model. |

#### S2. Normalization and Benchmark Structure

Raw Hamiltonian terms have different numerical scales and combinatorial growth with sequence length. Each contribution is therefore converted to a random-reference Z-score using non-designed sequences with the same length, alphabet, and environmental parameters as the evaluated sequence. Typical decoy sets contain 10,000–60,000 sequences depending on the calculation; the interfacial AMP recognition and one-vs-rest libraries use 3,000 random references per sequence because they require repeated penetration and environment scans. The web implementation allows the random-reference sample size to be configured.

The benchmarks are organized to test distinct failure modes. Homogeneous aqueous and apolar-core reference helices test whether the model assigns the expected environmental preference to experimentally characterized peptide classes. Canonical membrane-active helices test anionic-interface recognition against non-AMP helical controls. Candidate-generation experiments then use the same weighted Hamiltonian under cross-environment and one-vs-rest objectives, and landscape diagnostics test whether low-energy sequences form coherent amphipathic basins.

#### S3. Supplementary Evidence Map

The main-text claims are supported by the following supplementary material. Section S11 lists the empirical amino-acid parameters used by the Hamiltonian. Section S12 reports the term-level normalization figure, final term weights, and representative encoding resources. Section S13 provides supporting dictionary and QAOA workflow figures. Section S14 contains boundary-case design checks, interfacial wheel examples, additional basin diagnostics, complete cross-environment candidates, complete AMP recognition scores, and complete one-vs-rest anionic-specificity libraries.

#### S4. QAOA as a Quantum-Compatible Optimization Backend

In the main text, QAOA is treated as one optimization backend for exploring the same physically motivated design Hamiltonian that is also studied with classical solvers. The peptide design problem involves selecting a sequence of amino acids that minimizes a multi-term energy function encoding local propensities, pairwise interactions, solvation, electrostatics, and neighbor-sensitive structural couplings. Formally, this can be represented by a classical Hamiltonian $H_C$ acting on a set of $n$ qubits, where each qubit encodes a binary variable corresponding to a bit in the position-specific amino acid code.

QAOA provides a natural framework for exploring this type of discrete physical model because:

- It constructs a variational quantum state that encodes superpositions of all possible amino acid sequences, allowing exploration of exponentially large configuration spaces.

- Alternating between the problem Hamiltonian $H_C$ and a mixing Hamiltonian $H_M$ guides the quantum state toward low-energy basins of the sequence landscape.
- The algorithm acts as a statistical sampler, shifting the output distribution toward an ensemble of highly probable sequence candidates rather than deterministically converging to a single point.

#### S5. QAOA Ansatz and Hamiltonians

This section provides a detailed derivation of the QAOA circuit and Hamiltonians, expanding upon the overview given in the main text.

The QAOA ansatz prepares a variational state by alternating unitary evolutions under two non-commuting Hamiltonians:

1. **Problem Hamiltonian $H_C$:** Encodes the classical energy function of the peptide sequence. In this implementation, $H_C$ is diagonal in the computational basis (Pauli-$Z$), with each term representing contributions from amino acid properties (hydrophobicity, charge, helix propensity), pairwise interactions (MJ energies), solvent/interface exposure, electrostatics, and neighbor-dependent helical couplings. The transverse hydrophobic moment is evaluated downstream as an amphipathicity descriptor and alignment rule, not as an explicit energetic contribution in the final Hamiltonian. The ground state of $H_C$ corresponds to the lowest-energy sequence under the specified environment and objective.

2. **Mixing Hamiltonian $H_M$:** Promotes transitions between basis states to explore the configuration space. A transverse-field operator is used,

$$H_M = \sum_{j=1}^{n} \sigma_j^x, \tag{1}$$

   whose ground state is the uniform superposition $|+\rangle^{\otimes n}$, enabling non-trivial exploration of sequence space.

For a circuit depth of $p$ layers, the variational ansatz is

$$|\psi(\vec{\beta}, \vec{\gamma})\rangle = \prod_{k=1}^{p} \left( e^{-i\beta_k H_M} \, e^{-i\gamma_k H_C} \right) |+\rangle^{\otimes n}, \tag{2}$$

with variational parameters $\vec{\beta}, \vec{\gamma} \in \mathbb{R}^p$, optimized using a classical routine.

#### S6. Problem-Inspired Parameter Initialization

To improve convergence and reduce sensitivity to local minima, a linear ramp initialization inspired by quantum annealing is employed:

$$\beta_k^{(0)} = \pi \left( 1 - \frac{k-1}{p} \right), \tag{3}$$

$$\gamma_k^{(0)} = \pi \frac{k-1}{p}, \tag{4}$$

for $k = 1, \ldots, p$. This schedule ensures that early layers primarily apply mixer-driven rotations, keeping the state close to uniform superposition, while later layers progressively emphasize the problem Hamiltonian $H_C$, enhancing energy separation between favorable and unfavorable sequences. The upper bound of $\pi$ for $\beta_1^{(0)}$ reflects a full Bloch sphere rotation under $H_M$, which creates near-maximal superposition; the lower bound of 0 for $\gamma_1^{(0)}$ ensures that the first layer applies minimal problem-Hamiltonian weighting, preserving the initial uniform exploration of configuration space.

#### S7. Classical Optimization Using COBYLA

The variational parameters $(\vec{\beta}, \vec{\gamma})$ are optimized with the gradient-free *Constrained Optimization BY Linear Approximation* (COBYLA) algorithm. COBYLA iteratively constructs local linear approximations of the objective function and adapts the trust region at each step. Its gradient-free nature is particularly advantageous for variational quantum algorithms, where finite-shot sampling introduces stochastic noise in energy estimates.

To mitigate non-convexity, multiple independent optimization runs are performed. The first run uses the linear ramp initialization, while subsequent runs are initialized randomly to enhance the probability of reaching the global minimum.

#### S8. Risk-Averse Objective Function (CVaR)

Instead of minimizing the average energy $\langle H_C \rangle$, a *Conditional Value at Risk* (CVaR) objective is adopted for small systems. CVaR focuses on the lowest-energy outcomes, reflecting the practical goal of identifying a single optimal sequence:

$$C_{\text{CVaR}} = \frac{1}{\lceil \alpha K \rceil} \sum_{j=1}^{\lceil \alpha K \rceil} E_j, \tag{5}$$

where $\{E_1, \ldots, E_K\}$ are the sorted energies from $K$ measurements, and $\alpha = 0.1$ selects the lowest 10% of energies. Minimizing CVaR increases the likelihood that the sampled sequence corresponds to the ground state. For larger systems, the standard expectation value is minimized for numerical stability.

#### S9. Final Solution Selection

After optimization, the QAOA circuit is executed with a high number of shots to estimate the output probability distribution. The algorithm does not deterministically yield a single sequence, but rather generates a statistical ensemble. The final chosen sequence candidates are extracted as the highest-probability modes of this distribution and then rescored with the same Hamiltonian used in the main text. This procedure identifies the most favorable configurations sampled from the energy landscape while keeping the biological interpretation attached to the Hamiltonian rather than to the quantum optimizer itself.

The choice of quantum-circuit evaluation method depends critically on the entanglement structure of the ansatz and the size of the system. While Tensor Network methods like Matrix Product States (MPS) offer polynomial scaling for weakly entangled 1D systems, the highly connected nature of the peptide design Hamiltonian and the entanglement generated by the QAOA ansatz typically require large bond dimensions to maintain accuracy. Consequently, exact Statevector evaluation was used for small instances ($L \leq 6$) to guarantee precision. For larger instances, a hardware-efficient heuristic strategy was adopted by reducing circuit depth and shot counts, rather than relying on approximate MPS methods that might degrade due to entanglement entropy growth.

#### S10. Dimensionality Reduction for Extended Sequences

To optimize longer sequences such as $L = 8$ with the quantum-compatible backend, the full 16-amino acid dictionary ($\log_2(16) = 4$ qubits/residue) demands 32 qubits, exceeding the practical simulation budget used here. To circumvent this, the size of the amino acid dictionary $\mathcal{A}$ was reduced to 8 residues. This reduction requires only $\log_2(8) = 3$ qubits per position, thereby lowering the total system requirement to 24 qubits and allowing the QAOA workflow to execute successfully.

A $k$-means clustering algorithm was applied to the 3D property space with the number of clusters defined as $k = 8$ (Fig.S1 A). Once the algorithm converged, the 3D centroid for each of the 8 clusters was calculated. To ensure that the algorithm evaluated real molecular entities, the representative amino acid for each cluster was selected by finding the residue with the minimum Euclidean distance to the computed cluster centroid in the 3D space.

For the optimization of even longer sequences ($L = 12$), a further reduction of the amino acid dictionary $\mathcal{A}$ was necessary to keep the quantum-compatible instances within the same practical simulation budget. By limiting the dictionary to 4 residues, the requirement drops to $\log_2(4) = 2$ qubits per position. As a result, the total system requirement is maintained at 24 qubits ($12 \times 2$), ensuring the problem remains computationally tractable. Following the same geometric and mathematical rationale, the $k$-means algorithm was re-initialized with $k = 5$ (subsequently discarding proline to yield a minimal 4-amino acid dictionary). The continuous 3D space was partitioned into distinct physicochemical regions, and the single closest actual amino acid to each of the new centroids was extracted to form the minimal dictionary (Fig.S1 B).

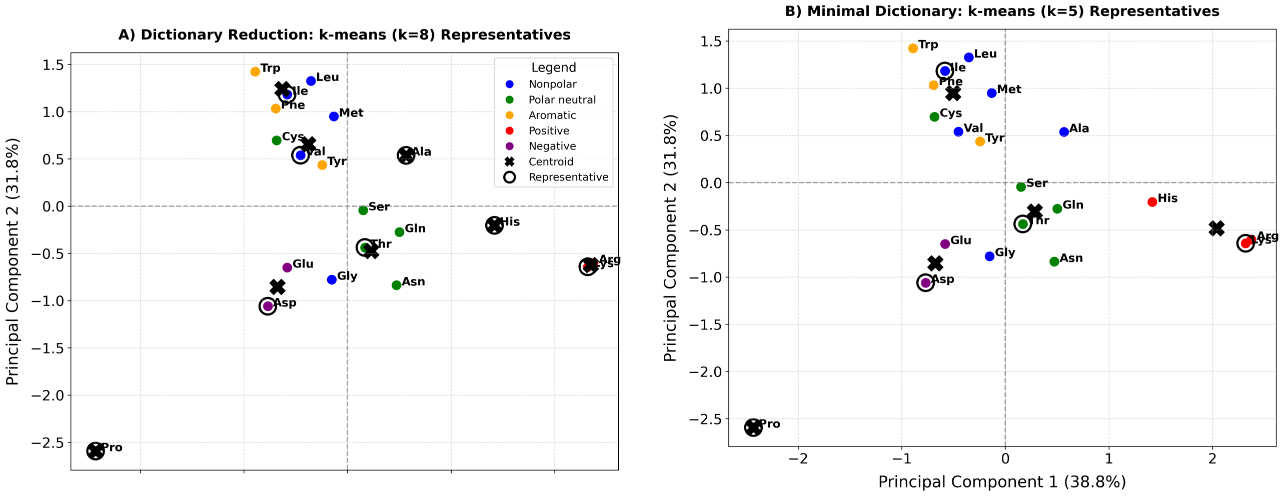


Figure S1: **Dimensionality reduction of the amino acid dictionary via $k$-means clustering.** Principal Component Analysis (PCA) projection of the 20 standard amino acids based on their normalized physicochemical properties (hydrophobicity, helix propensity, and charge). The axes represent the first two principal components, which together capture 70.6% of the total dataset variance. Amino acids are color-coded according to their predefined physicochemical classes. **(A)** Selection of the 8-residue dictionary ($|\mathcal{A}| = 8$). Black crosses indicate the mathematical cluster centroids determined by the $k$-means algorithm ($k = 8$) computed in the full 3D property space. Hollow black circles highlight the selected representative amino acids, identified by minimizing the Euclidean distance to each respective centroid. **(B)** Selection of a minimal 4-residue dictionary ($|\mathcal{A}| = 4$) using $k = 5$ clusters, with Proline subsequently excluded due to its helix-breaking properties, following the same geometric rationale. This demonstrates the adaptive partitioning of the physicochemical space to accommodate varying hardware constraints in the quantum algorithm.

### S11. Empirical Amino Acid Parameters

| Residue | Helix propensity ($h_\alpha$) (kcal/mol) |
|---|---|
| Ala | 0.00 |
| Leu | 0.21 |
| Arg | 0.21 |
| Met | 0.24 |
| Lys | 0.26 |
| Gln | 0.39 |
| Glu | 0.40 |
| Ile | 0.41 |
| Trp | 0.49 |
| Ser | 0.50 |
| Tyr | 0.53 |
| Phe | 0.54 |
| Val | 0.61 |
| His | 0.61 |
| Asn | 0.65 |
| Thr | 0.66 |
| Cys | 0.68 |
| Asp | 0.69 |
| Gly | 1.00 |
| Pro | 3.15 |

Table S2: Helix propensities of the 20 amino acids (in kcal/mol) [1].

| Amino Acid | $H_\alpha$ | Type |
|---|---|---|
| Asp (D) | −0.77 | Charged (−) |
| Glu (E) | −0.64 | Charged (−) |
| Lys (K) | −0.99 | Charged (+) |
| Arg (R) | −1.01 | Charged (+) |
| His (H) | 0.13 | Titratable (neutral here) |
| Gly (G) | 0.00 | Nonpolar |
| Ala (A) | 0.31 | Nonpolar |
| Val (V) | 1.22 | Nonpolar |
| Leu (L) | 1.70 | Nonpolar |
| Ile (I) | 1.80 | Nonpolar |
| Pro (P) | 0.72 | Nonpolar |
| Met (M) | 1.23 | Nonpolar |
| Phe (F) | 1.79 | Aromatic |
| Trp (W) | 2.25 | Aromatic |
| Tyr (Y) | 0.96 | Aromatic |
| Thr (T) | −0.04 | Polar |
| Ser (S) | 0.26 | Polar |
| Cys (C) | 1.54 | Polar |
| Asn (N) | −0.60 | Polar |
| Gln (Q) | −0.22 | Polar |

Table S3: Fauchère–Pliska hydrophobicity scale [2].

#### S12. Hamiltonian Normalization, Weights, and Encoding Resources

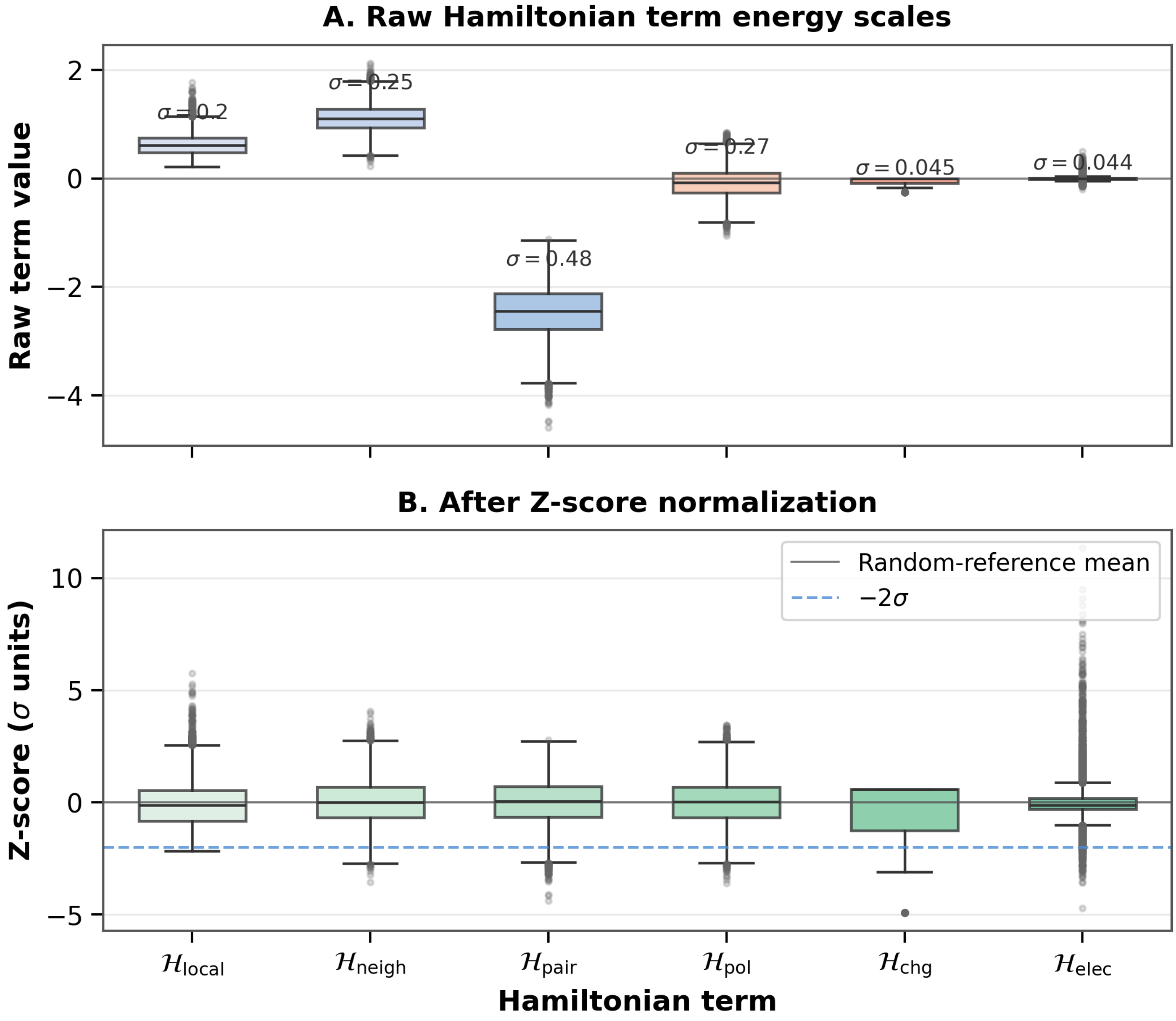


Figure S2: **Thermodynamic scaling and Z-score normalization of the Hamiltonian terms.** For a representative interfacial $L = 12$ setup, the six energetic Hamiltonian terms used in the model have heterogeneous raw scales before normalization and become directly comparable after Z-score standardization over $10{,}000$ random decoy sequences generated with the same length, alphabet, and environmental parameters.

Table S4: **Effective Hamiltonian weights used in the main analyses.** The transverse hydrophobic moment is retained only as a diagnostic and alignment descriptor, not as an energetic term.

| Component ($\mathcal{H}_i$) | Weight ($\lambda_i$) |
|---|---|
| $\mathcal{H}_{\text{chg}}$ | 3 |
| $\mathcal{H}_{\text{pol}}$ | 6 |
| $\mathcal{H}_{\text{elec}}$ | 4.0 |
| $\mathcal{H}_{\text{local}}$ | 1.5 |
| $\mathcal{H}_{\text{pair}}$ | 0.1 |
| $\mathcal{H}_{\text{neigh}}$ | 1.5 |

Table S5: **Representative qubit requirements for one-hot and compact binary encodings.**

| $L$ | $\lvert\mathcal{A}\rvert$ | One-hot | Binary | Reduction |
|---|---|---|---|---|
| 5 | 4 | 20 | 10 | 2× |
| 10 | 8 | 80 | 30 | 2.67× |
| 10 | 20 | 200 | 50 | 4× |

#### S13. Supplementary Methodological Figures

To support the reduced-alphabet construction used in the larger sequence calculations, the amino acids were examined in a normalized physicochemical space defined by hydrophobicity, helix propensity, and charge. The figures below complement the dimensionality-reduction discussion in the main text.

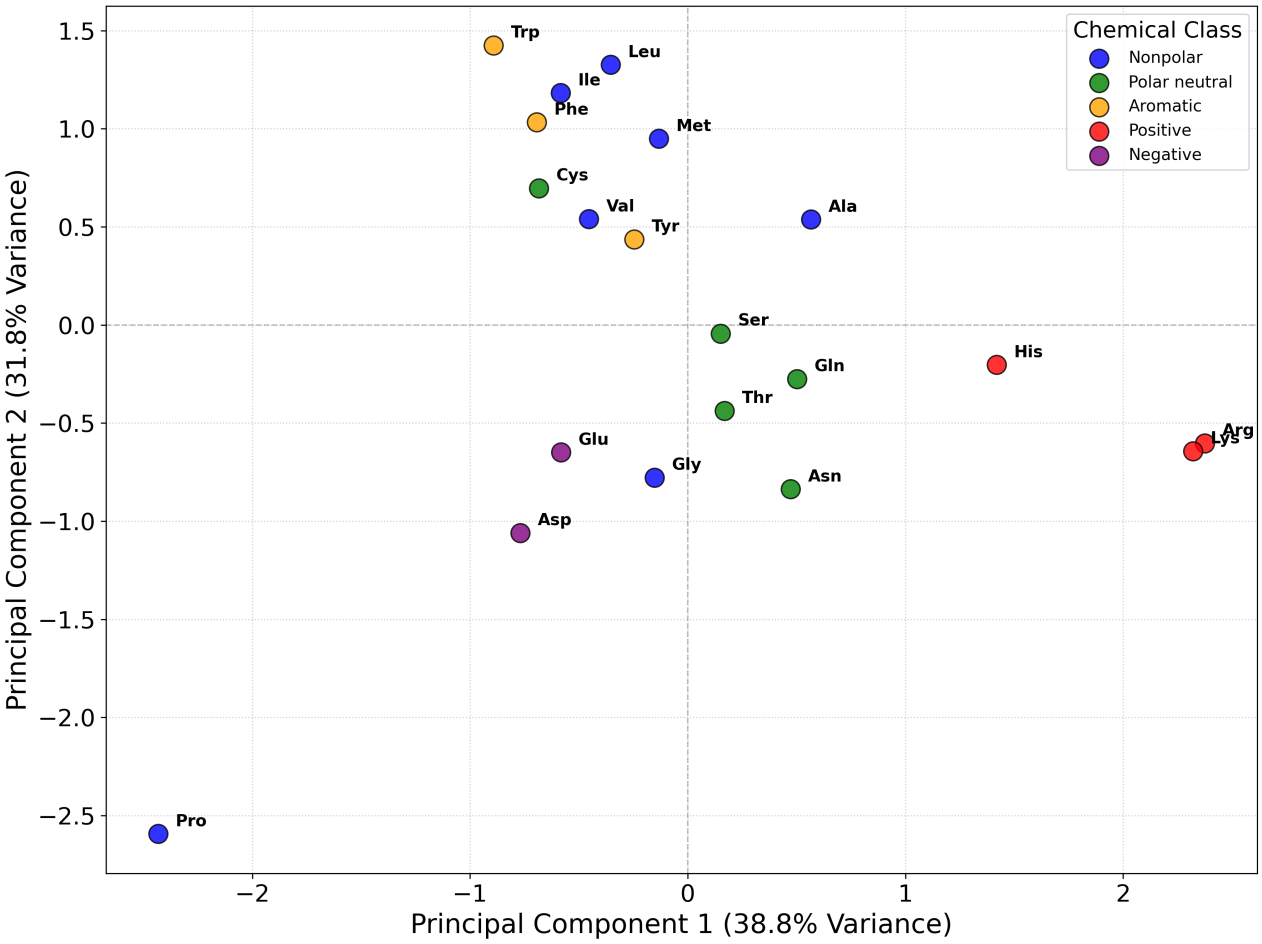


Figure S3: **Amino-acid similarity map via PCA.** Projection of the 20 amino acids onto the first two principal components, which together account for 70.6% of the total variance. Colors represent distinct chemical classes, highlighting clustering between residues with related physicochemical roles and the marked divergence of Proline due to its unique structural constraints.

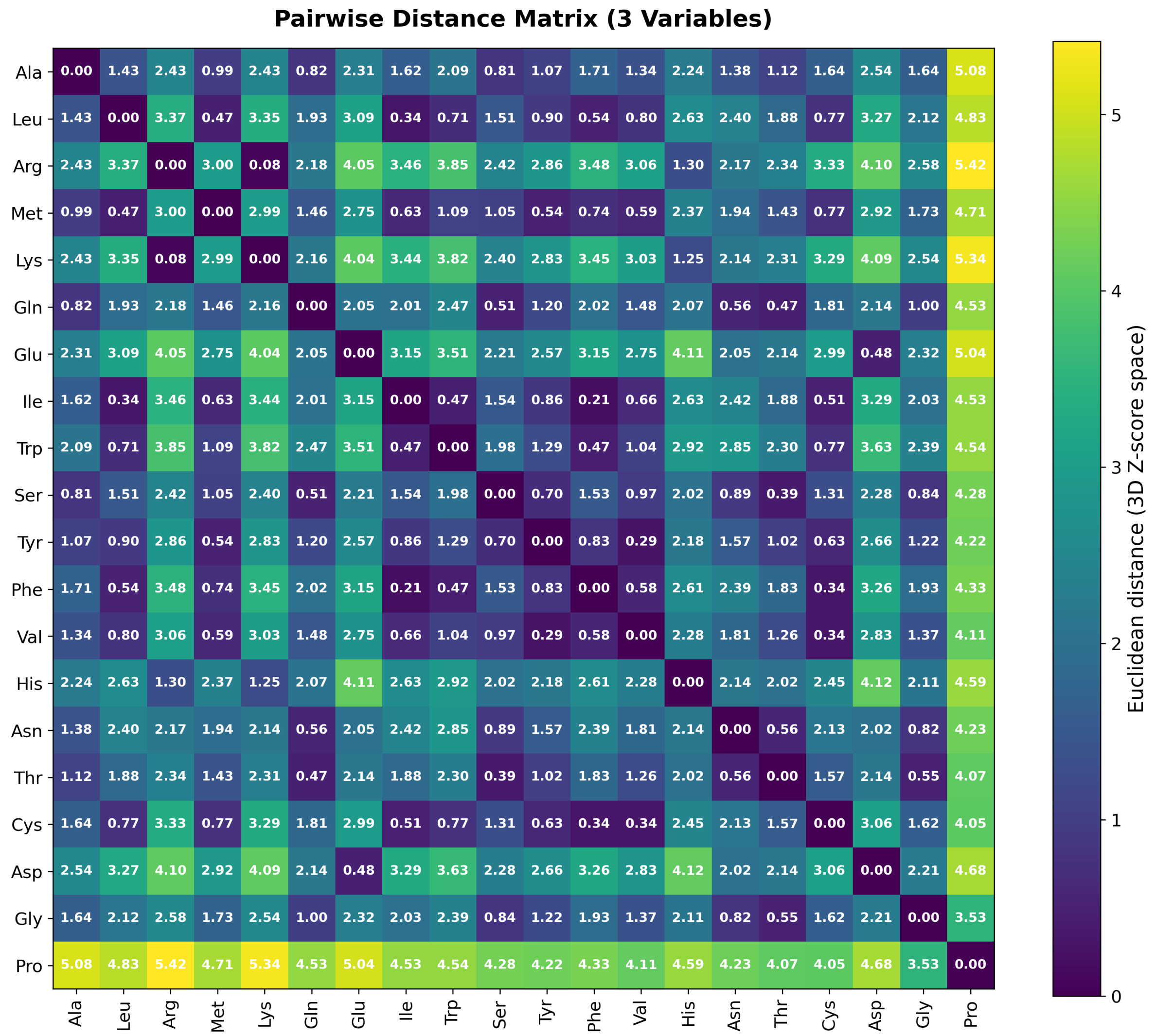


Figure S4: **Pairwise Euclidean distance matrix in the normalized 3D property space.** Darker shades indicate high biochemical similarity, while lighter shades emphasize strong physicochemical separation. The matrix provides a complementary quantitative view of the redundancy structure used to motivate reduced amino-acid dictionaries.

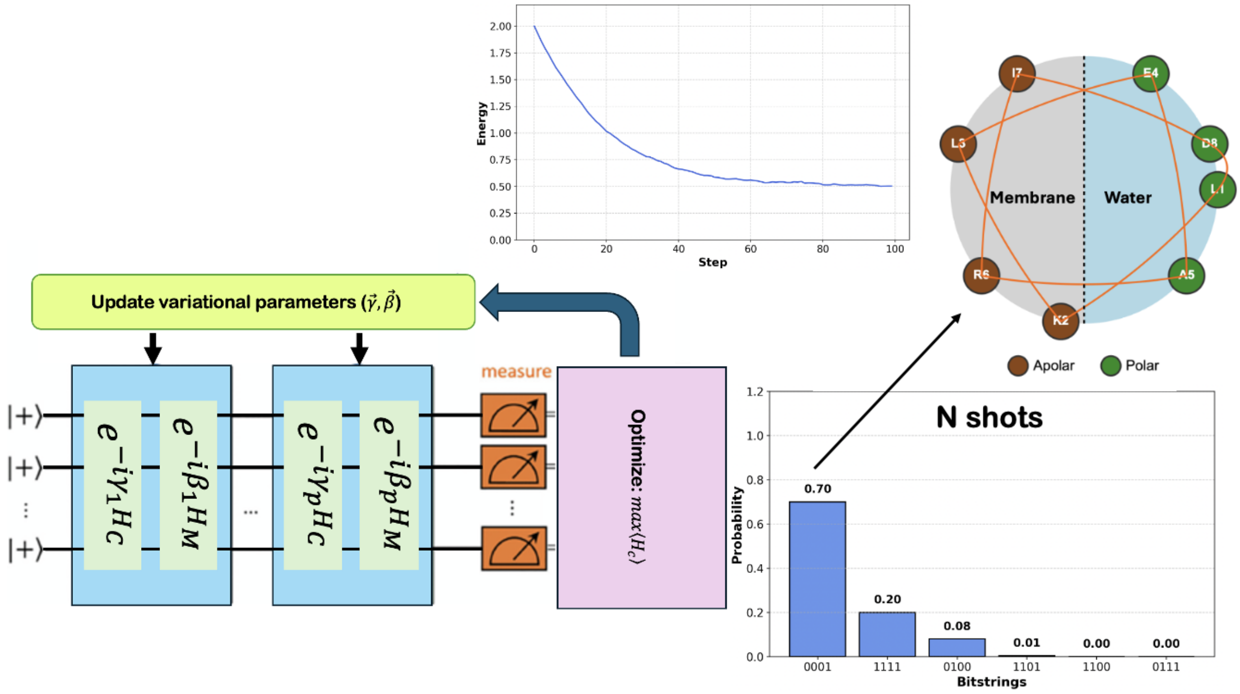


Figure S5: **Schematic representation of the QAOA workflow.** The variational circuit alternates between the problem Hamiltonian and a mixer Hamiltonian, with classical optimization of the circuit parameters between evaluation rounds.

#### S14. Supplementary Result Figures and Tables

The following figures provide result details that support, but are not essential to, the main narrative of the manuscript.

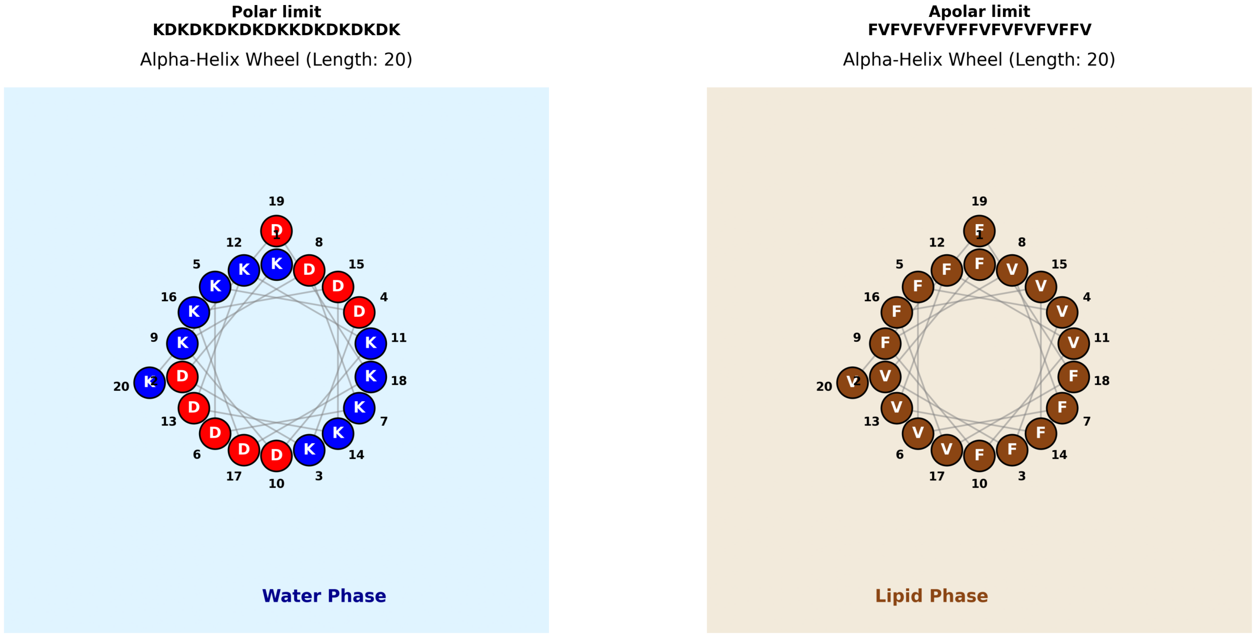


Figure S6: **Illustrative boundary-case solutions in fully polar and fully apolar environments.** The two panels show representative homogeneous-limit helices obtained from the same design framework in the polar and apolar limits. These examples are included as qualitative sanity checks on the directional force balance of the Hamiltonian and should be interpreted as thermodynamic boundary cases rather than as experimentally validated reference helices.

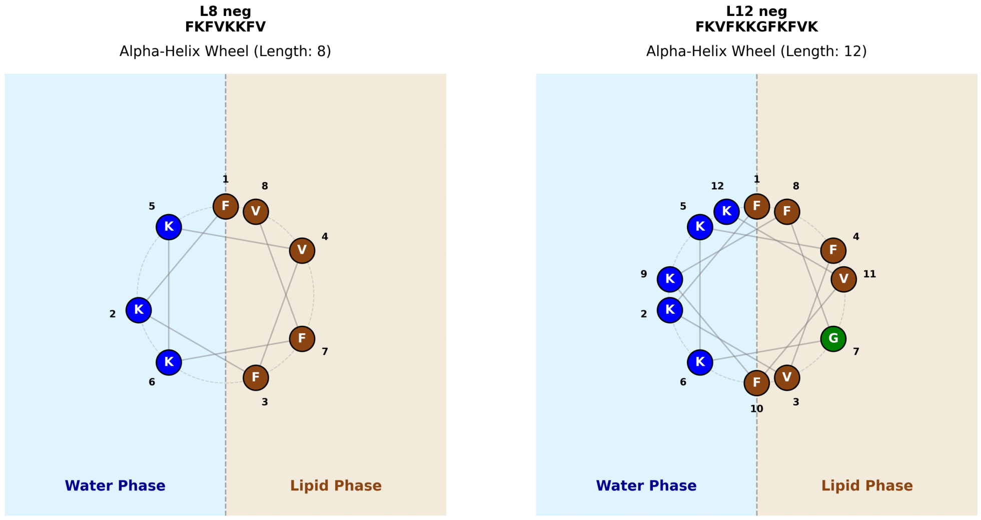


Figure S7: **Representative interfacial designs for short and intermediate helical scaffolds.** The panels show negative-interface candidate helices obtained with the same Hamiltonian at $L = 8$ and $L = 12$. These examples are included as qualitative wheel-level checks on amphipathic organization and charge placement; the main-text validation relies on reference-sequence recognition and Z-score benchmarks.

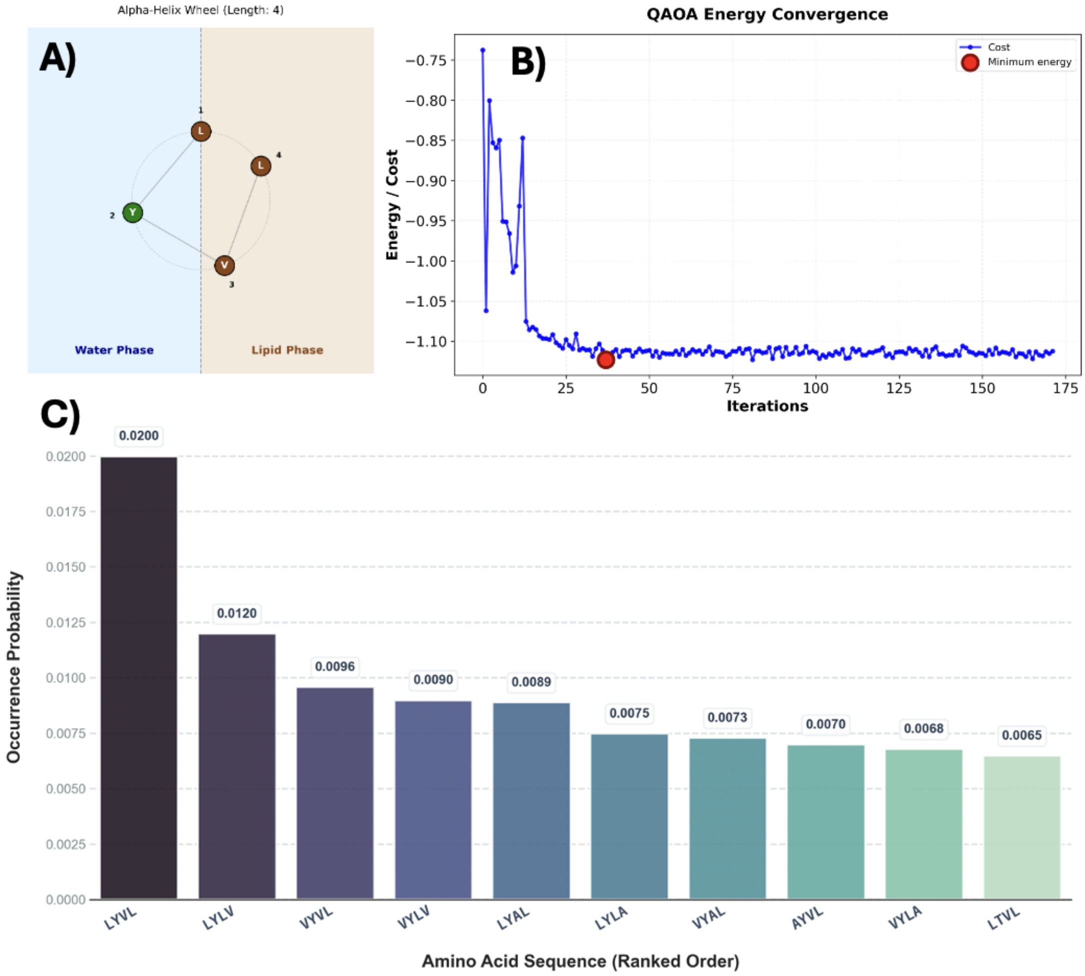


Figure S8: **Detailed QAOA output for the representative** $L = 4$ **interfacial design case. (A)** Alpha-helix wheel projection of one dominant low-energy sequence (LYVL). **(B)** Energy convergence of the QAOA objective during optimization. **(C)** Probability distribution over the top-ranked sequences sampled from the final variational state.

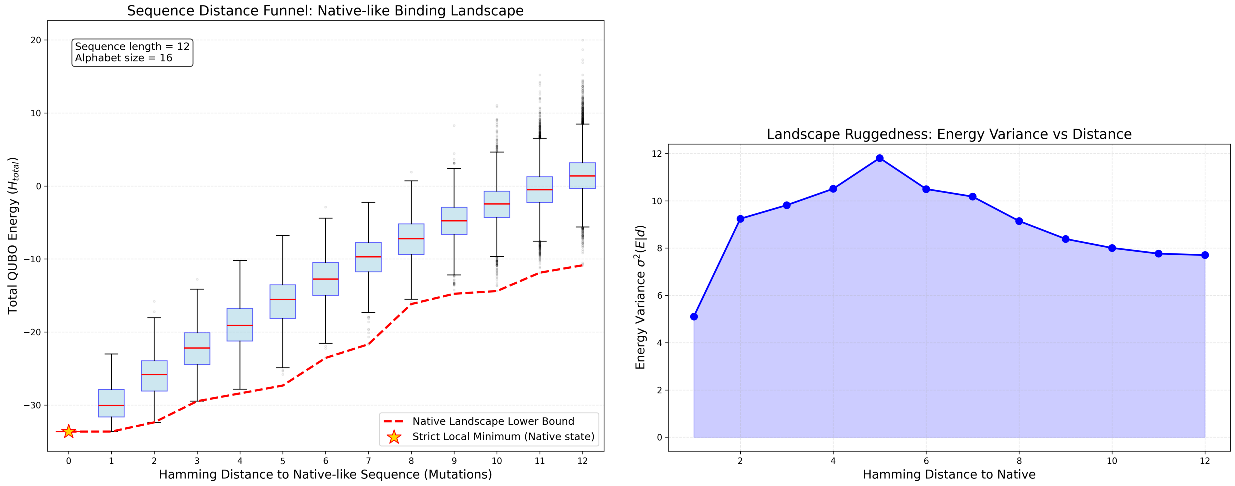


Figure S9: **Additional low-energy basin diagnostics. (Left)** Total Hamiltonian energy versus Hamming distance relative to a strict local minimum identified by steepest descent. The lower bound decreases as sequences approach the reference optimum. **(Right)** Conditional energy variance $\sigma^2(E|d)$ across sequences at fixed distance, highlighting maximal ruggedness at intermediate distances and a narrower variance near the low-energy basin.

Table S6: **Cross-environment-compatible helices obtained by minimizing** $\mathcal{H}_{\text{polar}} + \mathcal{H}_{\text{apolar}} + \lambda_{\text{gap}}|\mathcal{H}_{\text{polar}} - \mathcal{H}_{\text{apolar}}|$. The optimization uses the neutral-histidine parametrization described in the main text. Design Z-scores are reported in units of the random-reference standard deviation $\sigma$ relative to 10,000 random sequences of the same length and alphabet in each homogeneous environment.

| $\lambda_{\text{gap}}$ | Sequence | Residue types present | $Z_{\text{polar}}$ ($\sigma$) | $Z_{\text{apolar}}$ ($\sigma$) | $\lvert\Delta H\rvert$ |
|---|---|---|---|---|---|
| 0 | EKEKFEKEKEKEKFEKEKEK | E, F, K | -9.26 | -1.29 | 59.70 |
| 0.5 | EKEKWFWFKEKEWWWFWKEK | E, F, K, W | -3.78 | -3.43 | 0.46 |
| 1 | WWWFWWWFKEKEKFEKEKEK | E, F, K, W | -3.72 | -3.38 | 0.55 |
| 2 | EKWWFDKEKEWWFFWKEKEW | D, E, F, K, W | -3.60 | -3.34 | 0.08 |
| 4 | MKEWVFFWWEKWEVFKEKEK | E, F, K, M, V, W | -3.28 | -3.02 | 0.02 |

Table S7: **Interfacial recognition benchmark for known anionic-membrane-active helices.** For each peptide, the helical-wheel phase was aligned so that the transverse hydrophobic moment points toward the non-polar side, and the interfacial score was minimized over sampled penetrations from 5% to 95% in 5% increments. All Z-scores are reported relative to 3,000 random sequences of the same length over the full 20-amino-acid alphabet.

| Peptide | $L$ | $Z_{\text{water}}$ | $Z_{\text{apolar}}$ | $Z^*_{\text{anionic}}$ | $p^*_{\text{anionic}}$ | $Z^*_{\text{neutral}}$ | $p^*_{\text{neutral}}$ | $Z^*_{\text{cationic}}$ | $p^*_{\text{cationic}}$ |
|---|---|---|---|---|---|---|---|---|---|
| LL-37 | 37 | -2.16 | 0.61 | -5.51 | 10% | -4.49 | 10% | -4.36 | 10% |
| Melittin | 26 | 1.67 | 1.46 | -1.71 | 60% | -1.17 | 60% | -0.66 | 60% |
| Magainin 2 | 23 | -0.36 | 0.28 | -3.81 | 35% | -3.53 | 30% | -3.01 | 30% |
| Cecropin A | 37 | -1.12 | 0.68 | -2.60 | 25% | -2.00 | 25% | -1.67 | 25% |
| PGLa | 21 | -0.26 | 0.53 | -3.56 | 60% | -2.59 | 60% | -1.91 | 60% |
| Piscidin 1 | 22 | 0.28 | -0.73 | -3.98 | 95% | -3.50 | 35% | -2.67 | 35% |
| BP100 | 11 | 2.32 | 2.55 | -5.56 | 50% | -3.77 | 40% | -3.00 | 40% |
| (EAAAK)3 | 15 | -2.84 | 1.34 | -1.71 | 10% | -1.68 | 5% | -1.79 | 5% |
| Poly-Ala 15 | 15 | 0.49 | 1.68 | 1.07 | 10% | 0.86 | 5% | 1.10 | 5% |

Table S8: **Complete ranked one-vs-rest anionic-specificity designs for the hard-selective limit $\lambda = 0$.** For each fixed penetration depth, up to five unique $L = 20$ sequences were retained after classical annealing with $\mathcal{L}_0 = H_{\text{anionic}} - \min(H_{\text{neutral}}, H_{\text{cationic}}, H_{\text{polar}}, H_{\text{apolar}})$. All Z-scores are reported in units of $\sigma$ relative to 3,000 random sequences of the same length over the full 20-amino-acid alphabet. In all reported cases, the most competitive off-target is the neutral interface, so the specificity margin is $\Delta Z = Z_{\text{neutral}} - Z_{\text{anionic}}$.

| Penetration | Rank | Sequence | $Z_{\text{anionic}}$ | $Z_{\text{neutral}}$ | $Z_{\text{cationic}}$ | $Z_{\text{water}}$ | $Z_{\text{apolar}}$ | $\Delta Z$ |
|---|---|---|---|---|---|---|---|---|
| 10% | 1 | KRRIRRAKKKEKRCFKKEKR | -7.34 | -4.36 | -3.20 | 3.97 | 10.71 | 2.98 |
| 10% | 2 | RRRIKRVKRREKKLVKKDKK | -7.31 | -4.58 | -3.55 | 4.33 | 10.63 | 2.74 |
| 10% | 3 | KKRCKKWRRKDKRIHRRDRR | -7.31 | -4.57 | -3.43 | 4.02 | 10.57 | 2.75 |
| 10% | 4 | RRKCRKVRKREKKCCRKDKK | -7.12 | -4.09 | -3.03 | 4.57 | 10.64 | 3.03 |
| 10% | 5 | RRKLKKHKRREKKLCKKDRR | -7.22 | -4.29 | -3.07 | 3.73 | 10.68 | 2.94 |
| 25% | 1 | KRRIRRAKKKEKRCFKKEKR | -7.57 | -4.64 | -3.54 | 3.91 | 10.10 | 2.93 |
| 25% | 2 | RRRIKRVKRREKKLVKKDKK | -6.43 | -3.54 | -2.46 | 3.93 | 10.75 | 2.89 |
| 25% | 3 | KKRCKKWRRKDKRIHRRDRR | -6.21 | -2.95 | -2.06 | 4.23 | 10.77 | 3.26 |
| 25% | 4 | RRKCRKVRKREKKCCRKDKK | -6.14 | -2.92 | -2.22 | 4.17 | 10.66 | 3.22 |
| 25% | 5 | RRKLKKHKRREKKLCKKDRR | -7.61 | -4.58 | -3.56 | 3.84 | 10.69 | 3.03 |
| 50% | 1 | HRMMRKLDKLVRRIDKRCDK | -5.48 | -3.87 | -3.68 | -1.14 | 2.50 | 1.61 |
| 50% | 2 | WKVWRKMMRFIRRLIKKLFR | -6.02 | -3.75 | -2.94 | 4.49 | 4.18 | 2.27 |
| 50% | 3 | KKGLRRCRKPWKKFWKKFHK | -5.01 | -2.45 | -1.59 | 5.19 | 8.04 | 2.56 |
| 50% | 4 | WKFKKKANRYIRRQAKRGTK | -3.28 | -0.59 | -0.08 | 3.50 | 7.60 | 2.69 |
| 50% | 5 | VKDYRRLVKKVKKYYRKPAK | -4.75 | -1.99 | -1.31 | 2.83 | 6.39 | 2.76 |
| 75% | 1 | AKAGKRFIREYQKANRIWTK | -5.03 | -2.46 | -1.95 | -0.56 | 2.48 | 2.57 |
| 75% | 2 | HRQEKKWSREMWRKNKYIRK | -4.08 | -1.54 | -1.02 | -0.33 | 3.80 | 2.54 |
| 75% | 3 | TRVTKKIERSDSRMEKWLSK | -3.89 | -2.54 | -2.00 | -2.60 | 0.95 | 1.29 |
| 75% | 4 | HKDYRRQVRRWWKPIRHRNR | -2.54 | -0.27 | 0.07 | 1.31 | 5.06 | 2.27 |
| 75% | 5 | GRKFKRWIKMLSRLARAMDR | -4.71 | -2.31 | -1.71 | 1.50 | 3.51 | 2.40 |
| 90% | 1 | TRFEYIPSRCMFKKVRETTR | -4.13 | -1.98 | -1.58 | -0.83 | 1.23 | 2.14 |
| 90% | 2 | FKCMFTGARSINKCYKKCTR | -2.70 | -0.97 | -0.59 | 0.86 | 2.09 | 1.73 |
| 90% | 3 | KKALLHFERPVGKQMRIMIR | -4.40 | -2.44 | -1.89 | -0.12 | 0.93 | 1.96 |
| 90% | 4 | AKLTEQRWRQIHRRFRFLWK | -2.73 | -0.90 | -0.53 | 1.03 | 2.37 | 1.83 |
| 90% | 5 | FKLPFPMQKQNEKQLRWSKK | -2.45 | -0.70 | -0.35 | -0.18 | 1.74 | 1.75 |

Table S9: **Complete ranked one-vs-rest anionic-specificity designs for the representative positive-$\lambda$ branch ($\lambda = 0.50$).** For each fixed penetration depth, up to five unique $L = 20$ sequences were retained after classical annealing with $\mathcal{L}_\lambda = H_{\text{anionic}} + (\lambda - 1)\min(H_{\text{neutral}}, H_{\text{cationic}}, H_{\text{polar}}, H_{\text{apolar}})$. This $\lambda = 0.50$ table is shown as representative of the $\lambda > 0$ branch because the $\lambda = 0.25$ and $\lambda = 0.75$ libraries are nearly identical. All Z-scores are reported in units of $\sigma$ relative to 3,000 random sequences of the same length over the full 20-amino-acid alphabet. In all reported cases, the most competitive off-target is the neutral interface, so the specificity margin is $\Delta Z = Z_{\text{neutral}} - Z_{\text{anionic}}$.

| Penetration | Rank | Sequence | $Z_{\text{anionic}}$ | $Z_{\text{neutral}}$ | $Z_{\text{cationic}}$ | $Z_{\text{water}}$ | $Z_{\text{apolar}}$ | $\Delta Z$ |
|---|---|---|---|---|---|---|---|---|
| 10% | 1 | KKKEKKEKKKEKKEKKKDKK | -10.03 | -7.79 | -6.27 | -3.08 | 6.98 | 2.24 |
| 10% | 2 | KKKEKKEKKKEKKEKKKEKK | -9.72 | -7.53 | -6.46 | -2.86 | 6.94 | 2.19 |
| 10% | 3 | KKKEKKDKKKEKKEKKKDKK | -9.84 | -7.70 | -6.49 | -3.27 | 6.69 | 2.15 |
| 10% | 4 | KKKEKKEKKKEKKDKKKEKK | -10.10 | -7.73 | -6.16 | -2.94 | 6.84 | 2.37 |
| 10% | 5 | KKKEKKDKKKEKKEKKKEKK | -10.06 | -7.34 | -6.25 | -3.07 | 6.81 | 2.71 |
| 25% | 1 | KKKEKKEKKKEKKEKKKDKK | -9.63 | -7.10 | -5.71 | -3.23 | 6.59 | 2.53 |
| 25% | 2 | KKKEKKEKKKEKKEKKKEKK | -9.59 | -7.16 | -5.77 | -3.08 | 7.03 | 2.43 |
| 25% | 3 | KKKEKKDKKKEKKEKKKDKK | -9.74 | -7.12 | -5.56 | -3.24 | 6.76 | 2.62 |
| 25% | 4 | KKKEKKEKKKEKKDKKKEKK | -10.40 | -7.81 | -6.54 | -3.16 | 6.74 | 2.59 |
| 25% | 5 | KKKEKKDKKKEKKEKKKEKK | -9.77 | -6.93 | -5.82 | -3.17 | 6.83 | 2.85 |
| 50% | 1 | WKWWKKWEKWWKKWWKKEWK | -8.60 | -6.81 | -5.60 | 0.91 | 0.69 | 1.79 |
| 50% | 2 | WKWDKKWWKWWKKEWKKWWK | -8.13 | -6.22 | -5.06 | 0.73 | 0.67 | 1.91 |
| 50% | 3 | WKWEKKWWKWWKKEWKKWWK | -8.21 | -6.26 | -4.92 | 0.81 | 0.68 | 1.95 |
| 50% | 4 | WKWDKKWFKWWKKEWKKWWK | -8.28 | -6.17 | -5.00 | 0.70 | 0.55 | 2.10 |
| 50% | 5 | WKWEKKWFKWWKKEWKKWWK | -8.00 | -6.21 | -4.93 | 0.64 | 0.61 | 1.79 |
| 75% | 1 | WKWWKKWWKWWWKWWKWWWK | -7.83 | -5.79 | -5.13 | 3.62 | -1.08 | 2.04 |
| 75% | 2 | WKWWKKWEKWWWKWWKWWWK | -8.60 | -7.01 | -5.64 | 1.82 | -1.72 | 1.59 |
| 75% | 3 | WKWWKKEWKWWWKWWKWWWK | -6.82 | -5.36 | -4.61 | 1.74 | -1.71 | 1.46 |
| 75% | 4 | WKWWKKWEKWWFKWWKWWWK | -8.57 | -6.61 | -5.50 | 1.64 | -1.62 | 1.95 |
| 75% | 5 | WKWWKKWWKWWEKWWKWWWK | -8.43 | -6.67 | -5.48 | 1.89 | -1.64 | 1.75 |
| 90% | 1 | WKWWWWWWKWWWKWWKWWWK | -7.61 | -6.10 | -5.31 | 3.41 | -3.59 | 1.51 |
| 90% | 2 | WKWWWWWWKWWWKWWKFWWK | -6.84 | -5.60 | -4.68 | 3.21 | -3.44 | 1.24 |
| 90% | 3 | WKWWFWWWKWWWKWWKWWWK | -8.32 | -6.99 | -5.99 | 3.35 | -3.55 | 1.33 |
| 90% | 4 | WKWWWFWWKWWWKWWKWWWK | -7.03 | -5.59 | -4.75 | 3.33 | -3.40 | 1.44 |
| 90% | 5 | WKWWWWWWKWWFKWWKWWWK | -8.28 | -6.86 | -5.84 | 3.22 | -3.49 | 1.42 |

**Near-invariance across the positive-$\lambda$ branch.** Across $\lambda = 0.25$, 0.5, and 0.75, the top-ranked candidate at each penetration depth is unchanged. The corresponding top-five libraries are also nearly identical, so only the complete $\lambda = 0.50$ table is shown here as representative of the positive-$\lambda$ branch. The only visible difference in the present calculations is a minor swap in the order of two near-degenerate candidates at 90% penetration for $\lambda = 0.75$ relative to $\lambda = 0.25$ and 0.5. Thus, once the absolute target term carries any nonzero weight, the one-vs-rest design branch becomes highly stable to further reweighting.
